# Multiplex spatial systems analysis of local nanodose drug responses predicts effective treatment combinations of immunotherapies and targeted agents in mammary carcinoma

**DOI:** 10.1101/2021.09.01.458631

**Authors:** Zuzana Tatarova, Dylan C. Blumberg, James E. Korkola, Laura M. Heiser, John L. Muschler, Pepper J. Schedin, Sebastian W. Ahn, Gordon B. Mills, Lisa M. Coussens, Oliver Jonas, Joe W. Gray

**Affiliations:** Department of Biomedical Engineering, OHSU Center for Spatial Systems Biomedicine; Knight Cancer Institute; Department of Cell and Developmental Biology Oregon Health & Science University, Portland, OR 97239, USA; Department of Radiology, Brigham & Women’s Hospital Harvard Medical School, Boston, MA 02115, USA; Division of Oncologic Sciences Oregon Health & Science University, Portland, OR 97239, USA

**Keywords:** personalized combination treatments, tumor microenvironment, implantable microdevice, spatial systems analyses

## Abstract

Better methods are needed to identify effective combinations of immunotherapies with chemotherapies and targeted anti-cancer agents. Here we present a Multiplex Implantable Microdevice Assay (MIMA) system for rapid *in vivo* assessment of the effects of multiple, spatially separate anticancer drugs directly in the complex tumor microenvironment. In prototypic experiments, olaparib, lenvatinib, palbociclib, venetoclax, panobinostat, doxorubicin, and paclitaxel and combinations thereof were administered simultaneously to murine mammary tumor models. Quantitative multiplex immunohistochemistry and spatial systems analyses of each local drug response defined cellular relations of fibroblasts, endothelial cells, immune lineages, immunogenic cell death, tumor proliferation and/or cancer stem cells that were used to predict effective drug combinations. A predicted combination of panobinostat, venetoclax and anti-CD40 showed long-term anti-tumor efficacy in multiple mouse models with no observable toxicity when administered systemically. Future MIMA use promises to design effective drug combinations for tumor cell control and immune activation on a personalized basis.

## INTRODUCTION

Modern cancer therapies increasingly seek to effect tumor control by simultaneously attacking tumor intrinsic vulnerabilities, enhancing anti-tumor immune activity and/or mitigating stromal mediators of resistance. Targeted drugs typically are designed to attack genetic or transcriptional vulnerabilities on which tumor cells depend for survival but non-malignant cells do not.

Genomic screening approaches have supported such tumor-intrinsic aspects of precision medicine, leading to matching of genomic aberrations with specific targeted agents (Li et al., 2021). In breast cancer, treatments targeting tumors that depend on estrogen receptor (ER) signaling, aberrant signaling resulting from human epidermal growth factor receptor 2 (HER2) amplification and/or over expression, CDK4/6 signaling and defects in DNA repair in triple negative breast cancer (TNBC) have been particularly effective (Bettaieb et al., 2017; Hanker et al., 2020; Harbeck et al., 2021; Lord and Ashworth, 2017). Unfortunately, these treatments are not uniformly effective even in primary tumors carrying the target and are usually only transiently effective in metastatic disease (Brady et al., 2017; Janiszewska et al., 2021; Jeselsohn et al., 2017). This may be due in part to drug modulation of aspects of the tumor microenvironment (TME) including immune function that negatively influence cancer control. This suggests that treatment efficacy can be increased by combing these drugs with agents that increase immunogenicity and/or counter microenvironment-mediated resistance, a hypothesis that we address in this paper.

The concept of enhancing cancer treatment efficacy by combining chemotherapies and targeted drugs with agents that enhance immune-mediated cancer cell killing is increasingly well established. The clearest example is the use of immunotherapies, including immune checkpoint blocking (ICB) antibodies as complements to tumor targeted therapies in various liquid and solid malignancies (Dummer et al., 2020; Robert, 2020). However, many cancers do not benefit from ICB including in breast cancer where efficacy has been limited to a subset of TNBC patients (Adams et al., 2019; Force et al., 2019). This lack of efficacy is attributed in part to two mechanisms: i) Low antigenicity through decreased expression of major histocompatibility complex class I (MHC-I)—observed mainly in luminal ER+ BC (Brady et al., 2017; Lee et al., 2016) and HER2+ BCs (Inoue et al., 2012; Janiszewska et al., 2021); and ii) a naturally immunosuppressive TME associated mainly with TNBC and HER2+ BC (Denardo et al., 2011; Gil Del Alcazar et al., 2017; Guerrouahen et al., 2020). Both of these mechanisms may limit the CD8+ T cell-mediated anti-tumor response, which then cannot be leveraged to improve efficacy of ICB therapies (Palucka and Coussens, 2016). Combinations of conventional chemotherapies and/or targeted anticancer drugs that increase immunogenic cell kill promise significant improvements in overall outcome (Galluzzi et al., 2018). However, further understanding of drug-immune system interactions is required to design effective and safe immune modulating combinatorial regimens.

A variety of experimental approaches have been deployed to elucidate the effects of drug combinations on the tumor and stromal components and to identify biomarkers that inform on the efficacy of treatment combination decisions (Letai, 2017). Biomarkers typically are identified by establishing associations between tumor features and outcomes in population-based clinical studies (Hellmann et al., 2018; Hugo et al., 2016) such as those supported by The Cancer Genome Atlas (Hutter and Zenklusen, 2018) and Human Tumor Atlas Network (Rozenblatt-Rosen et al., 2020) programs. However, these association-based approaches need to be tested for causality in systems that faithfully recreate the interactions of the various components of the TME. Common model systems include tumors that arise in immune competent mice and short- or long-term ex vivo cultures comprised of tumor and stromal components using miniscule scaffolds and active fluidics to closely model specific aspects of the TME (Deng et al., 2017; Jenkins et al., 2017; Tatárová et al., 2016). However, these studies in mice typically are slow, expensive and labor-intensive, and comprehensive modeling of tumor-microenvironment interactions in ex vivo systems remains a major challenge (Yuki et al., 2020).

We report now on a high-throughput in vivo approach to rapidly, safely and efficiently assess the effects of multiple drug combinations on TME composition and architecture in living mouse models that is also applicable to humans (Dominas et al., 2021). Our study focuses on mouse mammary cancers and our approach integrates two recently introduced high-throughput and high-content techniques: a minimally invasive implantable microdevice (IMD) (Jonas et al., 2016, 2015; Watson et al., 2018) for intratumor delivery of nanoliter doses (nanodoses) of multiple drugs or drug combinations into spatially separate regions, and multiplexed immunohistochemistry (mIHC) (Lin et al., 2015; Tsujikawa et al., 2017) to assess the in-situ responses of the tumor-microenvironment milieu near each drug delivery site. Computational analyses of serial mIHC staining and imaging of 30+ proteins allow comprehensive categorization of standard immune and non-immune stromal cell types and states as well as complementary characterization of tumor proliferation, apoptosis, differentiation and/or immunogenicity. Assessment of the composition and spatial distribution of the functionally different cell types in each drug delivery area facilitates identification of drug-mediated mechanisms of response and resistance that rapidly reveal new therapeutic intervention strategies. We refer to the overall approach as the Multiplex Implantable Microdevice Assay (MIMA).

We used the MIMA to evaluate the effects of FDA approved drugs olaparib, palbociclib, doxorubicin, venetoclax, panobinostat, lenvatinib and paclitaxel and combinations thereof on the tumor and TME, in the immunocompetent MMTV-PyMT (mouse mammary tumor virus-polyoma middle tumor-antigen) mouse model. This commonly used genetically engineered mouse model for breast cancer research mirrors many aspects of human breast cancer progression and heterogeneity (Attalla et al., 2021; Guy et al., 1992; Lin et al., 2003). Out of eight treatments tested, five showed unique, significantly enriched histopathological signature that we used to predict synergistic TME modulating combination treatments that subsequently were validated in traditional systemic dosing studies. Among these, the combination of panobinostat, venetoclax and agonist anti-CD40 monoclonal antibodies (mAB) provided the strongest response for long-term disease control in multiple models of mammary carcinoma and was well-tolerated.

## RESULTS

### MIMA components and design to identify effective TME modulating combination treatments

The IMD used for drug delivery in the MIMA system was a 5.5 mm long, 0.75 mm diameter biocompatible resin cylinder capable of delivering multiple drugs or drug combinations at up to 18 spatially separate regions inside a living tissue per device (Figure 1A). IMDs were loaded with drugs in pegylated semi-solid form within the wells of the cylinder so that drugs were released upon implantation via passive diffusion (Jonas et al., 2015). Local concentrations of drugs in the IMD were tuned to produce drug levels at each site in the tissue to match those achieved during systemic treatment in clinical practice (Figure S1A and Table S1) (Jonas et al., 2016). Importantly, the miniscule nanodoses of drugs delivered via IMDs do not generate the toxicities typically associated with systemic treatments (Jonas et al., 2015).

**Figure 1.**
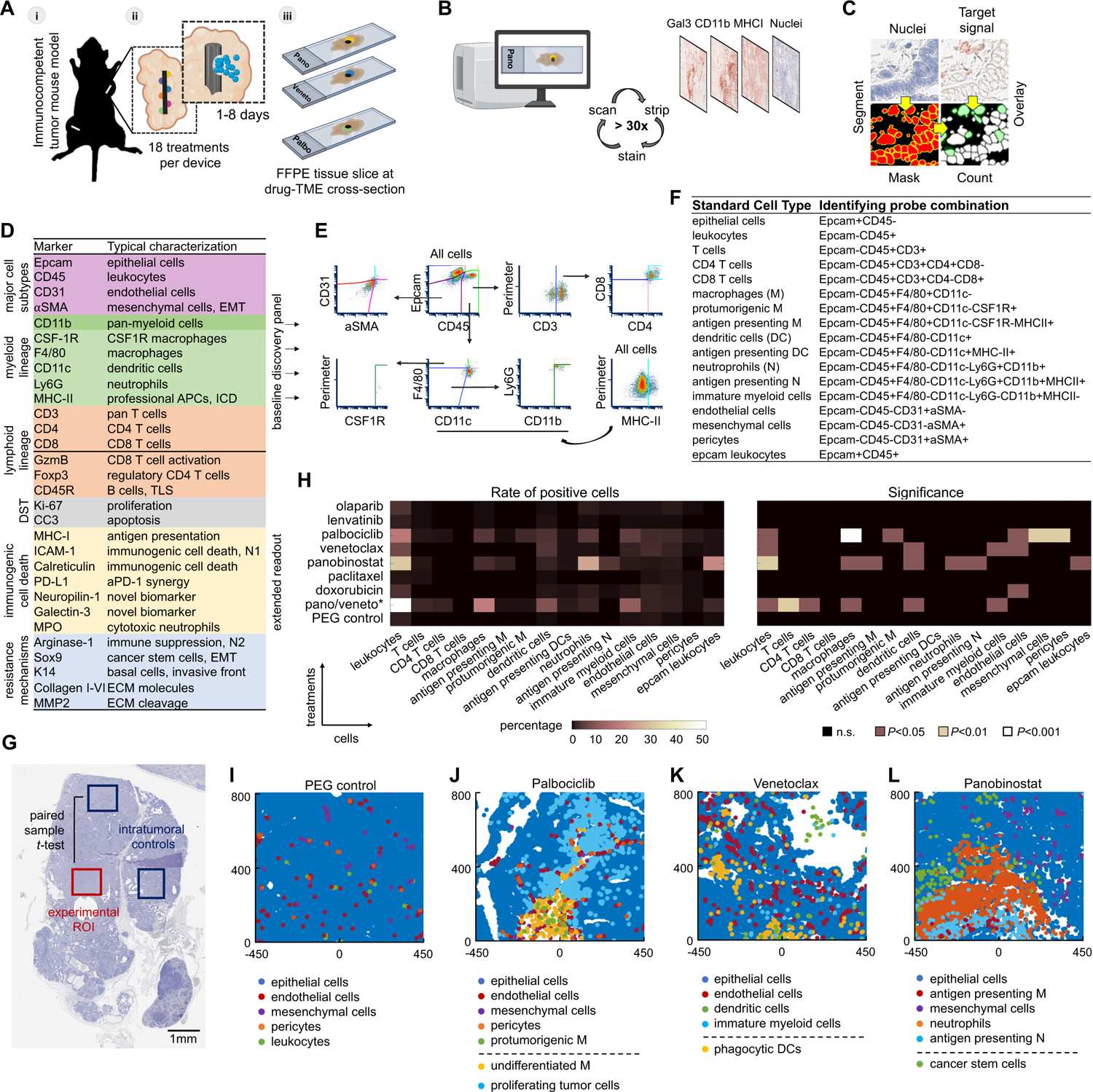
MIMA components and testing of locally induced drug effects on TME. (A) Schematic of IMDs implanted into a multifocal mouse model of mammary carcinoma (i). Treatments are released from device reservoirs into spatially separated regions of tumors through passive diffusion (ii). Following incubation, the tumor is extracted with the device in place and is formalin fixed paraffin embedded (FFPE). Each condition is assayed individually (iii) using 30+ color staining (B, D). (B) Schematic of the mIHC technique composed of iterative histological staining on a single FFPE slide which is alternated with digital scanning microscopy to detect the target marker. (C) Acquired images are co-registered with nuclear staining and the mean intensity of antibody staining within a mask is calculated for each cell to count marker positive cells in an intact tissue. (D) Antibody list used to interrogate a broad range of tumor intrinsic and tumor-microenvironmental states as categorized and color-coded in the table. DST; drug sensitivity testing. (E) Multidimensionality reduction in hierarchical gating to classify standard cell types. (F) List of probe combinations from the baseline discovery panel (D, top half) identifying standard cell types. (G) Macroscopic view of the hematoxylin-stained tumor tissue implanted with IMD. Experimental condition (red box, assay area) was compared to control, untreated intra-tumor, regions distant from the drug-releasing site (blue boxes). To obtain greater control over cofounding variables, paired sample one tailed t-tests were used to determine enrichment of induced TME states. (H) Heatmap of mean percentage of positive cells (left) and level of significance (right) at depicted targeted agents and chemotherapies (y-axis). Polyethylene glycol (PEG) served as negative control. Total cell counts to define percentage of positivity were between 3000 to 5000 cells per assay area and were matched ± 300 total cells for paired samples (experimental vs control region). Minimum population proportion within 5% margin of error and 95% confidence level was set to 0.75% (represents 12 cells) to discriminate noise from specific signal. n=3 wells from 3 tumors from 2-3 mice per treatment. MMTV-PyMT mice with late stage spontaneously growing tumors were implanted for three days. *For panobinostat/venetoclax (pano/veneto) condition details see Figure 6C. (I-L) Presentation of selected standard cell types in XY space. [0,0] coordinate is the drug releasing site; and direction of release is upward.

After treatment for 1 to 8 days, tumors were harvested with the IMD in place, prepared as formalin-fixed, paraffin-embedded (FFPE) samples and serial tissue sections were cut orthogonal to the axis of the IMD (Figure 1A). Sections through each drug delivery well in the IMD were analyzed implementing mIHC (Figure S1B) according to published procedures (Chang et al., 2017; Tsujikawa et al., 2017) to assess drug effects using a range of markers with specificity being cross-validated using cyclic immunofluorescence (cycIF) (Lin et al., 2015) (Figure S1C-F). mIHC generated multiprotein images of each tissue section through a process of serial immunostaining, imaging and stripping of each FFPE section (Figure 1B, Figure S1B) using antibodies to proteins that define cell types and/or functions *in situ* (Figure 1C, D and S2A). In our studies, 30+ proteins were interrogated for each section. Each mIHC image was analyzed by segmenting individual cells and calculating cell positivity for each segmented cell (Figure 1C, S2B-D). The mIHC antibody panel (Figure 1D, Table S2 and S3) was specifically designed to capture a broad range of TME states and to identify actionable phenotypes with preferential detection of early and local responses (Table S4).

We implemented a binary gating strategy (Figure 1E, S2E) using measurements of 13 proteins (Epcam, CD45, CD31, *α*SMA, CD3, CD4, CD8, CD11b, F4/80, CSF1R, CD11c, Ly6G, MHC-II; Figure 1D, S2A; baseline discovery panel) to define 17 major tumor cell types or states with focus mostly on immune and non-immune stromal cells. We refer to these cells as “standard cell types” hereafter. They included, for example, T cells (Epcam-CD45+CD3+), antigen presenting macrophages (Epcam-CD45+F4/80+CD11c-MHC-II+), immature myeloid cells (Epcam-CD45+F4/80-CD11c-Ly6G-CD11b+MHC-II-) and endothelial cells (Epcam-CD45-*α*SMA-CD31+). For the full list see Figure 1F. We measured the abundances and spatial organizations of these standard cell types for all test conditions to narrow down the target stromal phenotypes in a controlled and unbiased manner. Then, we interrogated additional proteins to refine the standard cell types and/or to report on other aspects of tumor and stromal cell biology including on basic drug sensitivity (CC3, Ki67), immunogenic cell death and/or processes typically associated with resistance or breast cancer progression such as cancer stem cells (Epcam+CD45-PyMT+Ki67-Sox9+), invasion (Keratin-14, K14); or immune suppression (arginase-1, arg-1) (Figure 1D; extended readout).

### Quantification of single cell events at local delivery sites reveal unique drug specific histopathological signatures of TME response

We used the MIMA system to perform a small-scale screening study in which we quantitatively assessed the responses to five FDA approved targeted therapies and two chemotherapeutic agents with distinct modes of action. The targeted drug were the poly (adenosine diphosphate [ADP]) ribose polymerase (PARP) inhibitor, olaparib (Lord and Ashworth, 2017); the multi-kinase vascular endothelial growth factor receptor (VEGFR)-1/2/3 inhibitor, lenvatinib (Kato et al., 2019); the cyclin dependent kinase (CDK)-4/6 inhibitor, palbociclib (Harbeck et al., 2021); the B-cell lymphoma (BCL)-2 inhibitor, venetoclax (Montero and Letai, 2018); and the pan-histone-deacetylase (HDAC) inhibitor, panobinostat (Atadja, 2009). The chemotherapeutic drugs were the DNA-intercalating agent, doxorubicin and the microtubule poison, paclitaxel, which are often used in first line therapy for breast cancer (Cardoso et al., 2018; Kumar et al., 2018). We assessed responses in late stage MMTV-PyMT mice with spontaneously growing tumors that mirror the morphology and aspects of progression of human breast cancers (Guy et al., 1992).

These tumors initially express ER strongly but expression decreases as they progress to late-stage carcinoma (Lin et al., 2003). Gene expression profiling reveal that tumors cluster with the luminal B subtype at the stage used herein (Herschkowitz et al., 2007; Lin et al., 2003). We chose a spontaneous rather than transplanted tumor model to better account for all stages of immune-biology associated with de novo tumor progression (Hanahan and Coussens, 2012), including editing (Dunn et al., 2004).

IMDs loaded with individual agents were implanted in tumors for three days since our previous work indicated that TME responses were apparent by this time (Watson et al., 2018). Our analyses of harvested tumors focused on the cell and molecular compositions and organizations that were significantly enriched in regions close to the drug deliver sites compared to remote intratumoral controls (Figure 1G). Quantification of the 17 standard cell types following seven candidate drug exposures revealed unique drug-specific histopathological signatures of TME response that are summarized in Figure 1H with Figures 1I-L showing computed images of the most prominent response cell types after treatment. Lenvatinib and paclitaxel produced no detectable local TME changes; olaparib was associated with a modest increase in macrophage, neutrophil and fibroblast density, whereas doxorubicin induced a significant increase in vasculature (Figure 1H). Palbociclib, venetoclax, and panobinostat produced significant ecosystem changes in both the immune and non-immune stromal states and thus we studied and described these targeted anti-cancer agents in more detail in the following sections. Predicted effective combinations derived from these studies (Table S4) were then validated in whole animals implementing commonly used murine mammary cancer cell lines (E0771 and/or EMT6 models (Herschkowitz et al., 2007)) allografted into syngeneic immunocompetent hosts.

### Palbociclib induces enrichment of CSF1R+ macrophages associated with pericyte branching and de novo tumor proliferation

Intratumoral treatment with the CDK4/6 inhibitor, palbociclib, induced a significant accumulation of several stromal cell types into the assay area including leukocytes, endothelial cells, pericytes and mesenchymal cells (Figure 1H, J, Figure 2A, Figure S3A, B). Among leukocytes, colony-stimulating factor 1 receptor (CSF1R)-positive macrophages were the most enriched subset (Figure 1H, 2A) but only 9% were positive for class II major histocompatibility complex (MHC-II) (Figure S3C) indicating these macrophages are not professional antigen presenting cells (APCs) and may be protumorigenic (Kowal et al., 2019; Reis E Sousa, 2006). CSF1R+ cells, in general, were uniquely and significantly enriched by palbociclib (Figure 2A, B and S3A, D), however, the marker was not solely expressed on macrophages as defined by the standard cell type classification (Figure 1F). Extended mIHC analyses revealed that CSF1R also was expressed on epithelial cells, fibroblasts and endothelial cells (Figure 2C). The majority (46%) of CSF1R+ cells were positive for the F4/80 macrophage and CD11b pan-myeloid markers, however, the CD45 leukocyte marker was under the detection limit of IHC (Epcam-CD45-CD31-*α*SMA-F4/80+CD11b+CSF1R+; Figure 2C, S3B, D) indicating they might be less differentiated protumorigenic macrophages (Deszo et al., 2001; Norazmi et al., 1989; Sophie Mokas et al., 2009). Spatial analyses showed that while the CD45+ macrophages were localized to regions immediately proximal to the drug delivery well, CD45-less-differentiated macrophages were localized both proximally and more distally and in some regions were associated with the contractile pericytes (Epcam-CD45-*α*SMA-CD31+) (Bergers and Song, 2005) (Figure 2B-D). This spatial distribution is apparent in a profile plot (Figures 2D, S3B) which shows the relative abundance of cells at increasing distances from the drug delivery well.

**Figure 2.**
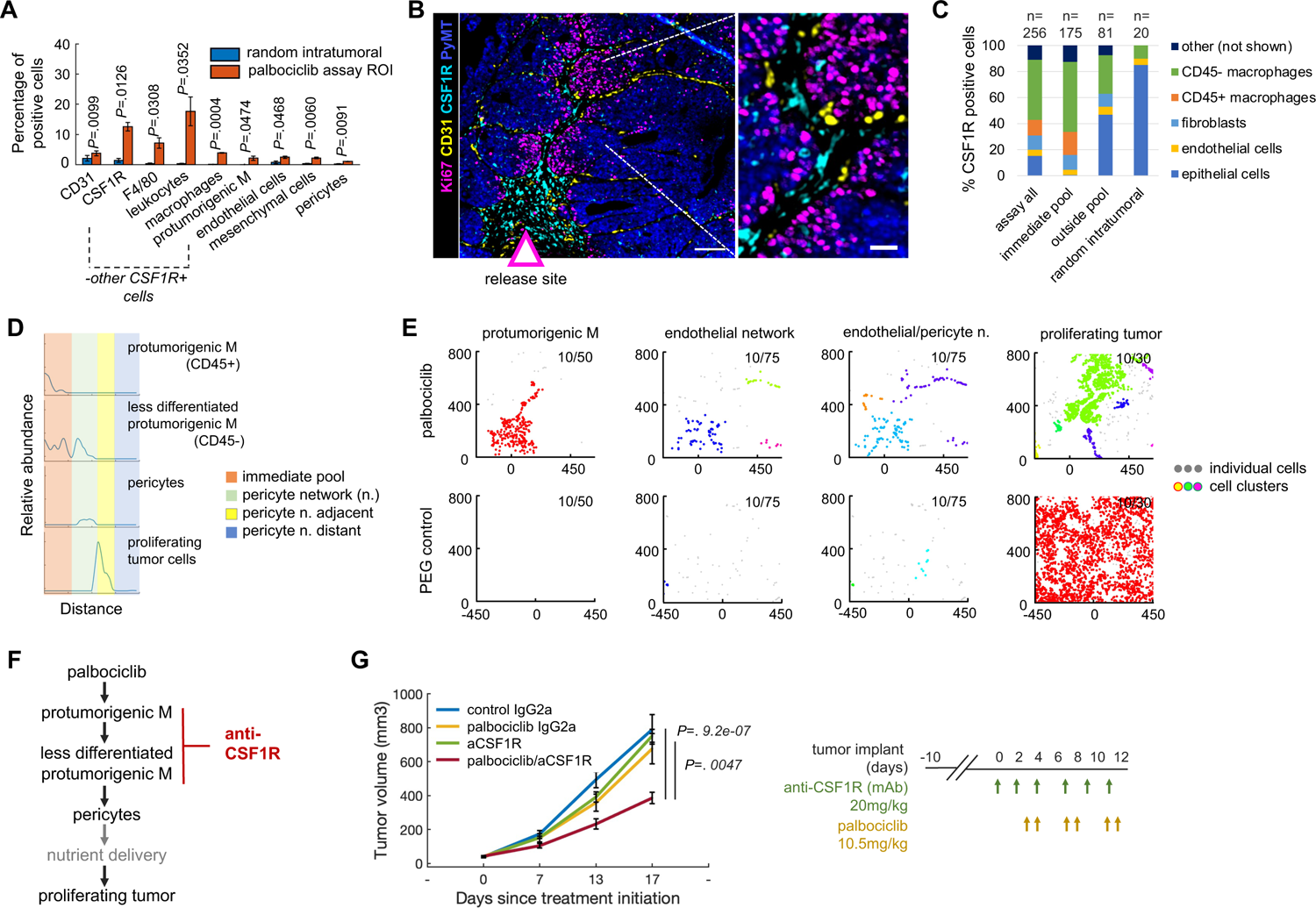
Local TME changes induced by targeted therapy palbociclib and whole animal studies testing the combination efficacy with predicted anti-CSF1R immunotherapy. (A) Quantification of single cell events induced by palbociclib using individual markers and standard cell type classification. Bars are mean ± s.e.; n=3 palbociclib reservoirs in three tumors from three different mice. Significance was calculated by paired sample one tailed t-test. Only significantly enriched cells are presented. For quantification of all TME lineages, see Figure S3A. (B) Sample composite image of key response markers. Arrow indicates the source and direction of palbociclib drug release. Dashed lines define the magnified area. Scale bar is 100μm (left); and 25μm (right). (C) Percentage of top five cell types expressing CSF1R stratified by zones in the palbociclib assay area. Immediate pool zone analyzed is visualized by the dashed line in Figure S3D. The number of cells (n) quantified is presented at the top of the figure. (D) Line profile of relative cell abundance as a function of distance from well (left to right). Assay zones are color-coded in the legend. (E) Distance-based clustering of depicted cell types as a set of XY coordinates. Coordinate [0,0] identifies the drug source. The direction of the drug release is upward. Clusters were identified by a minimum 10 cells within maximum distances of 50μm, 75μm and 30μm for CSF1R+ protumorigenic macrophages, endothelial/pericyte network and proliferating tumor cells, respectively. Each cluster is depicted with a randomized color; clusters were merged and share one color if present within the maximum distance range. Individual (non-clustering) cells are shown as smaller light gray points. (F) Palbociclib model of response presented as line diagram and site of intervention using immunotherapy depicted in red. (G) Tumor burden measurement of mice bearing EMT6 tumors after systemic treatment using drugs as color-coded in the graph. Shown is mean ± s.e.; n=8 to 10 tumors per group. Significance was calculated using an independent two-sample two-tailed t-test with equal variance.

We also assessed the propensity of specific cell types to cluster together by mapping the locations where 10 or more cells of a defined phenotype occurred together in regions 30, 50 or 75 μm in diameter (Figure 2E and S3E). The selection of cluster sizes was based on the highest variance between palbociclib treated and PEG control regions excluding strategies that showed treatment unspecific clusters in untreated regions (e.g. clusters of as few as 5 cells or distances of *≥*100 μm; Figure S3F). These analyses showed that the CSF1R+ macrophages and CD31+ endothelial cell/pericyte structures were organized together in response to palbociclib drug stimulus and did not appear in PEG control tissues. The patterns for the CD31+ cell aggregates were branch-like with pericytes integrated within endothelial structures suggestive of neovascularization and blood pressure/flow control (Bergers et al., 2003) (Figure 2E, S3E). The profile plot and distance-based cluster analyses also showed clusters of Ki67-positive neoplastic cells (Epcam-CD45+PyMT+Ki67+) distal to the drug delivery site and proximal to the macrophage-pericyte networks (Figure 2D, E and S3B, D and E) indicating that the macrophage-pericyte structures may be linked to the increased tumor cell proliferation in local microculture as summarized schematically in Figure 2F. The high density of CSF1R on multiple cell types and the associated increase in Ki67+ tumor cells suggested to us that compounds targeting the CSF1/CSF1R axis might enhance palbociclib efficacy by countering these CSF1R-mediated processes (Table S4).

We tested this concept in a systemic study of the EMT6 breast cancer model, by treating mice bearing orthotopically implanted tumors into the mammary fat pads of immunocompetent syngeneic mice with intraperitoneal injections of palbociclib, an anti-CSF1R antibody and a combination of the two. Whereas the individual drugs did not impact the rate of tumor growth, the combination treatment significantly reduced tumor growth rate (Figure 2G). Thus, the efficacy of palbociclib/anti-CSF1R revealed by analyses of responses to intratumoral treatments was affirmed in whole animal experiments.

### Venetoclax recruits phenotypically distinct clusters of dendritic cells, immature myeloid cells and endothelial cells

Intratumor treatment with the BCL-2 inhibitor venetoclax, did not produce significant cell death as indicated by IHC staining for cleaved caspase 3 (CC3) (see Figure 4C). However, it resulted in significant recruitment of leukocytes including dendritic cells (DCs), immature myeloid cells; and endothelial cells to the region proximal to the drug release well (Figures 1H, K and 3A and S4). Unlike palbociclib, the endothelial cells were not associated with *α*SMA (Figure 3B) suggesting they form small blood vessels (Bergers et al., 2003) that are not supported by pericytes. The recruitment of dendritic cells is noteworthy since they play important roles in regulating the balance between immune tolerance and activity (Domogalla et al., 2017).

**Figure 3.**
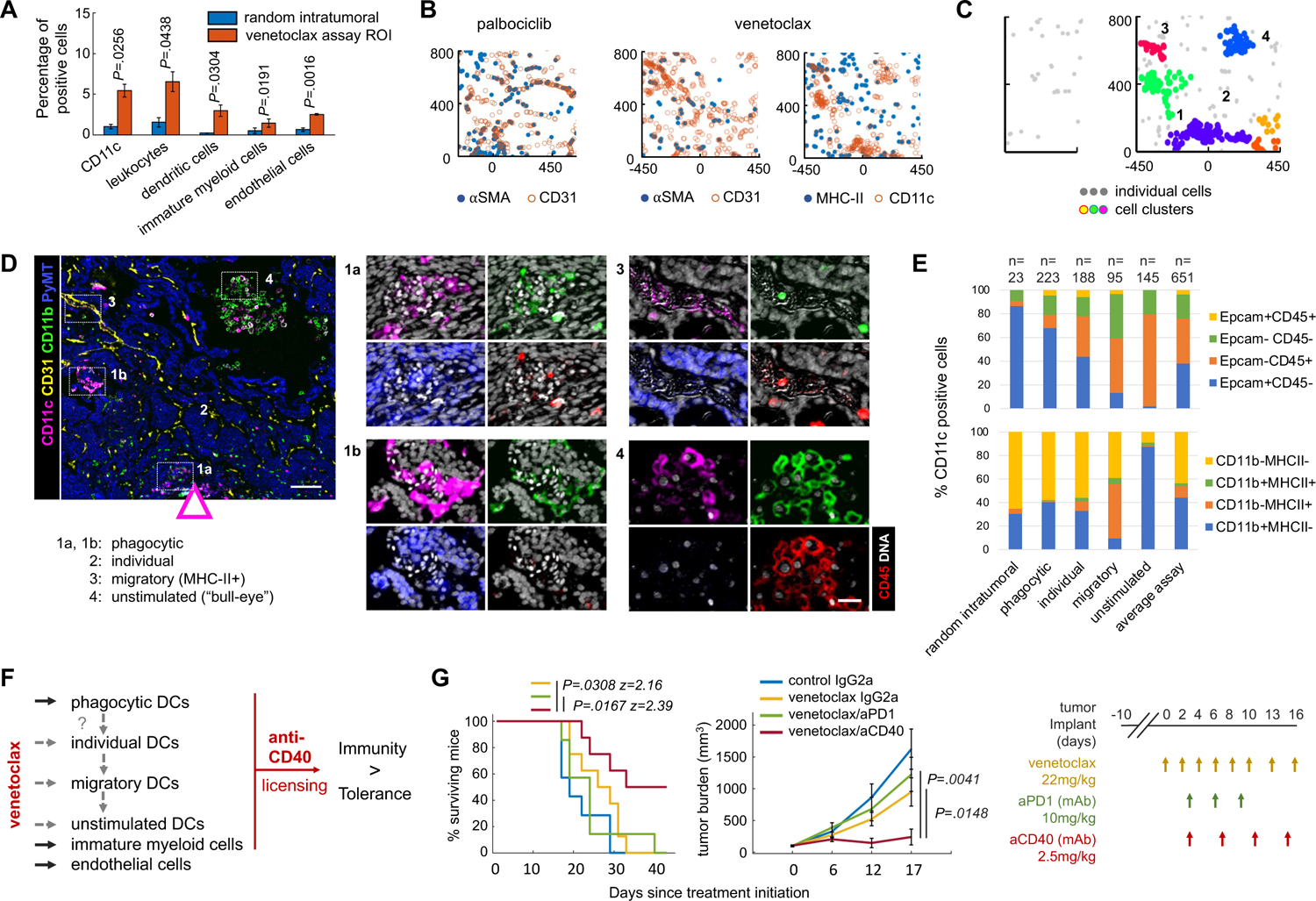
Local TME changes induced by Venetoclax and whole animal studies testing the combination treatment efficacy with the predicted anti-CD40 immunotherapy. (A) Quantification of single cell events using individual markers and standard cell types. Bars are mean ± s.e.; n=3 venetoclax reservoirs in two tumors from two different mice. Significance was calculated by paired sample one tailed t-test. Only significantly enriched cells are presented. For quantification of all cells, see Figure S4A. (B) Marker co-expression in XY coordinates in the palbociclib (left graph) and venetoclax (mid and right graph) assay area. Each color-coded dot represents a marker positive cell. Coordinate [0,0] identifies the drug source. The direction of the drug release is upward. (C) Distance-based cluster analysis of CD11c positive cells as a set of XY coordinates in random intratumoral (left) and venetoclax assay (right) regions. Clusters are displayed in randomized colors if at least 10 cells are present within maximum distance range 50μm; individual (non-clustering) cells are shown as light gray points. (D) Sample composite image of key response markers as color-coded on the left. The arrow indicates the source and direction of the venetoclax drug release. DHashed boxes define the magnified area on the right. Here individual markers are overlayed on DNA signal (in white). Scale bar 100μm; and 30μm for the magnified image. (E) Percentages of Epcam and CD45 (top) and CD11b and MHC-II (bottom) positive cells within morphologically different CD11c + cells presented as a stack bar graph. The number of cells (n) in each analysis is presented above the bar. Two to three ROIs from two venetoclax samples were analyzed and summed per each zone. (F) A model of venetoclax response presented as an influence diagram with sites of intervention using immunotherapy depicted in red. The relation of morphologically distinct and spatially separate CD11c DC clusters remains unclear (gray dashed arrows). (G) Survival rates (left) and tumor burden measurements (right) of mice bearing E0771 tumors after systemic treatment using drugs as color-coded in the line graphs. Shown is mean ± s.e.; n=7-8 mice per group. Significance was calculated by log-rank (Mantel-Cox) and by an unpaired two-tailed t-test with equal variance for survival and tumor burden rate, respectively. The treatment dose and schedule are presented schematically. For single treatment effects of anti-PD-1 and anti-CD40 see Figure 6H.

**Figure 4.**
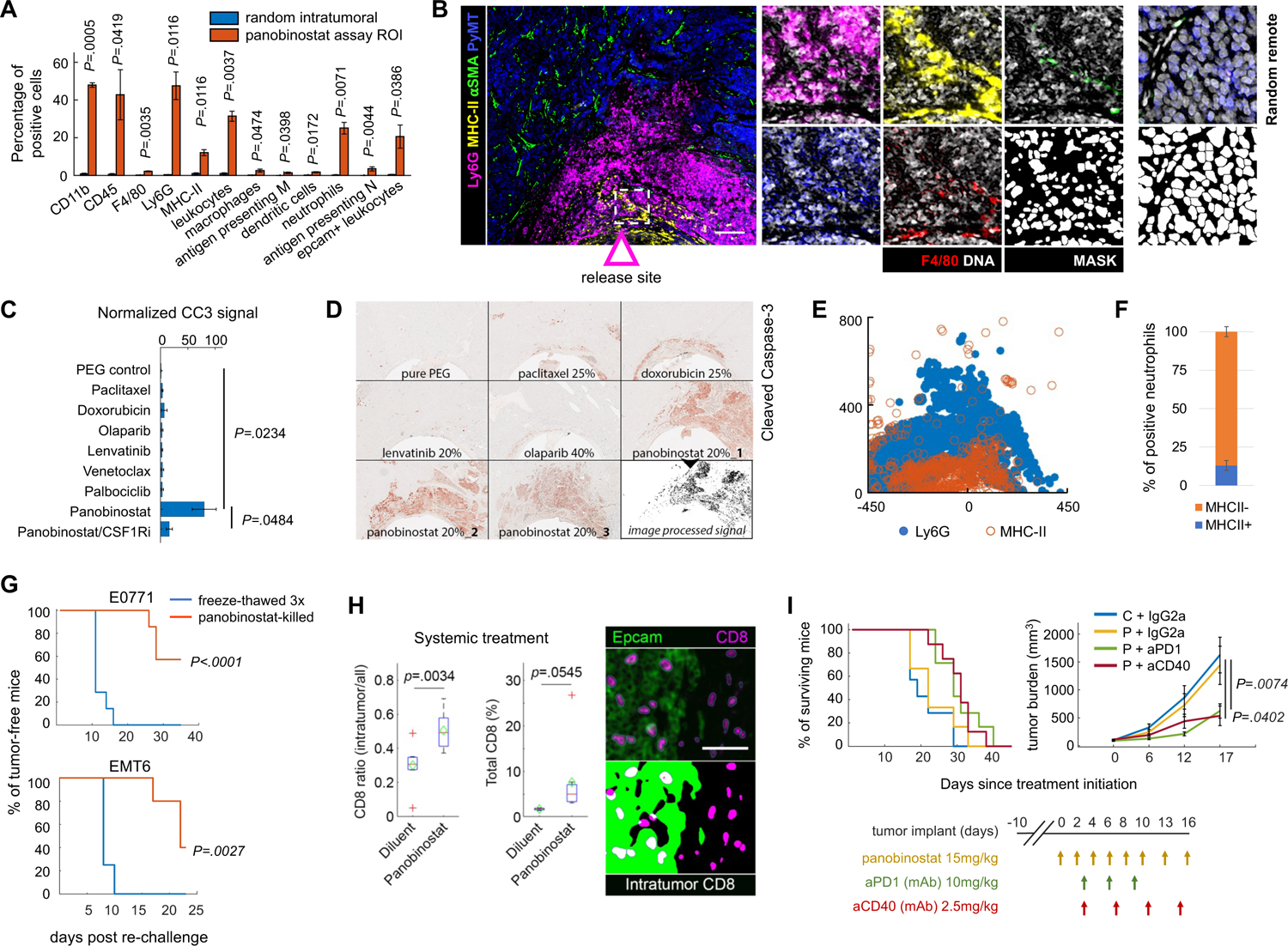
Local effects of panobinostat and whole animal studies testing induction of anti-tumor immunity in mouse mammary carcinoma. (A) Quantification of single cell events using individual markers and standard cell types. Bars are mean ± s.e.; n=3 panobinostat reservoirs in two tumors from two different mice. Significance was calculated by paired sample one tailed t-test. Only significantly enriched cells are presented here; for quantification of all cell, see Figure S5A. (B) Composite image of the most prominent markers appearing at the panobinostat reservoir as color-coded on the left. An arrow indicates the source and direction of the drug release. A dashed box defines the magnified area (right), which shows F4/80 staining in red and DNA signal and DNA-derived mask (white). Ly6G, MHC-II, aSMA or F4/80 expression was not enriched in random, drug-remote, intratumoral regions (far right). Scale bar, 100μm. (C) Quantification of PEG normalized average mean CC3 intensity (px value) in the assay region. The graph shows mean ± s.e signal intensity; n=3 wells from 3 tumors from 2-3 mice per treatment; significance was calculated using an independent two-sample t-test with equal variance. CSF1R inhibitor (PLX3397 in mouse chow at average 40mg/kg dose) was administered for seven days before the three-day-long IMD application. (D) CC3 IHC image of a sectioned tissue surrounding the IMD at depicted targeted agents and chemotherapies. Three replicates are presented for the most potent death-inducing drug, panobinostat. A computationally processed CC3 signal for the 20% panobinostat image is shown in the lower right as a binary image. (E) Marker co-expression in XY coordinates. Each dot represents a marker positive cell as color coded on the bottom. Coordinate [0,0] identifies the drug source. The direction of the drug release is upwards. (F) Percentage of MHC-II+ neutrophils. Shown is mean ± s.e.; n=3 panobinostat reservoirs. (G) Induction of anti-tumor immunity measured in a vaccination study using panobinostat treated cells and negative control (cells killed by three freeze/thaw cycles). Line graphs show percentages of mice free from palpable tumors. The P-value was calculated by log-rank (Mantel-Cox) test. n=7 per each group for E0771 model; and n=4 (control) and n=5 (experimental) for EMT6 model, respectively. (H) Quantification of intratumoral CD8+ T cell infiltration into ErbB2*Δ*Ex16 spontaneously growing tumors 7 days after systemic panobinostat treatment. The central mark indicates median and the bottom and top edges of the box indicate 25th and 75th percentiles, respectively. Green diamonds show the means. n=8 randomly selected ROIs using four tumors from two to three mice per group. A two-color composite image of Epcam and CD8 staining (top right) and a processed image showing intraepithelial CD8+ cells depicted in white (bottom right). Scale bar, 100μm. (I) Survival rates (left; 100% to 0%) and tumor burden measurements (right) of E0771 tumor bearing mice treated with depicted treatments as color-coded in the line graphs. Mean ± s.e. per timepoint are presented; n=8 to 12 mice per group. Significance was calculated by log-rank (Mantel-Cox) and by unpaired two-tailed t-test with equal variance for survival and tumor burden rate, respectively. The treatment dose and schedules are presented.

Interestingly, CD11c+ cells (primarily a dendritic cell marker) aggregated into clusters in regions near venetoclax delivery, but not in random intratumoral regions far from the drug releasing reservoir (Figure 3C). The clusters were phenotypically distinct as defined by their morphology and positivity for Epcam, CD45, MHC-II and CD11b (Figure 3D, E). The dendritic cell clusters located nearest to the drug delivery well were intermixed with neoplastic cells, exhibited brighter and smaller nuclei, (Figure 3D; 1a and1b) and greater than 60% of this population Epcam+CD45-(Figure 3E, S4C) suggesting these cells might be phagocytic DCs (Goodridge et al., 2011). Dendritic cells further from the reservoir were aligned and intermixed with endothelial cells (Figure 3D; 3), possibly resulting from migration from/to the TME and they were the only population expressing MHC-II (Figure 3B and E) (Kedl et al., 2017). Thus, only a small subset of the DCs recruited to the venetoclax well were antigen presenting cells. DCs distal from the reservoir were mostly present in tumor cleared areas (Figure 1K and 3D; top right). These cells were CD11b and CD45 positive (Figure 3E) and all three stains CD45, CD11b and CD11c showed a “bulls-eye” pattern around an unstained cytoplasmic area centered on the nucleus (Figure 3D; 4). The surface CD45 expression and the “bulls-eye morphology” suggest these cells were unstimulated DCs (Goodridge et al., 2011).

It is thus far unknown whether the spatially separated and phenotypically distinct DC clusters are functionally related, or whether they were induced by venetoclax as separate entities (Figure 3F). However, the literature suggests that MHC-II negative DCs have limited antigen presenting potential and limited capacity to prime T cells (Reis E Sousa, 2006). Additionally, unstimulated DCs as well as immature myeloid cells, which also were significantly enriched near the venetoclax well (Figure 1H, K, 3A, and S4A, C), have been reported to induce immunological tolerance (Domogalla et al., 2017; Griffiths et al., 2016; Sotomayor et al., 1999) raising the possible therapeutic utility of agents that could mitigate the tolerogenic potential of these cells (Table S4). Agonist monoclonal anti-CD40 antibodies act on DCs and immature myeloid cells by increasing their antigen presenting capacity, maturation and activation potential (called licensing) (Sotomayor et al., 1999). Licensed DCs have the capacity to shift the balance from tolerance to anti-tumor immunity (Griffiths et al., 2016; Hoves et al., 2018; Kowal et al., 2019). We reasoned that anti-CD40 immunotherapy could be used to enhance the anti-tumor immune capacity of DCs and immature myeloid cells that were recruited by venetoclax treatment (Figure 3F).

We tested this concept in E0771 breast cancer cells grown orthotopically in the mammary fat pads of immunocompetent C57BL/6 mice by treating systemically with intraperitoneal injections of venetoclax, an anti-CD40 agonist antibody and a combination of the two. Neither agent was effective as a single drug but the combination of the two significantly reduced the tumor growth rate and increased overall survival with 60% of mice surviving for >180 days (Figure 3G). For comparison, the combination of venetoclax with a programmed death ligand-1 (PD-1) inhibitory antibody did not significantly affect tumor growth rate or survival (Figure 3G). Again, a therapeutic strategy suggested by the MIMA proved to be effective in whole animal experiments.

### Panobinostat induces immunogenic cell death associated with recruitment of antigen presenting neutrophils and macrophages

Intratumor delivery of the pan-HDAC inhibitor, panobinostat, led to significant recruitment of several immune cell populations including dendritic cells, antigen presenting macrophages and (antigen presenting) neutrophils (Figure 1H, L, 4A and S5A, B). The cells located *immediately* at the drug releasing well were MHC-II positive antigen presenting macrophages (Figure 1L, 4B, 5A and S5B-C). The presence of such macrophages inside tumor nests has been associated with better clinical responses, improved survival and improved efficacy of PD-1/programmed death ligand-1 receptor (PD-L1) therapies (Guerriero et al., 2017; Kawai et al., 2008). Depletion of these macrophages using a CSF1R inhibitor significantly decreased panobinostat-induced cell death (Figure 4C) implying this immune subset has a functional, anti-tumor role. This result is in line with previous studies showing that concurrent depletion of macrophages abrogates the anti-tumor efficacy of HDAC inhibitors (Guerriero et al., 2017).

Neutrophils were the most abundant immune population recruited to the panobinostat well (Figure 1H, 4A) and these were organized in a band slightly farther from the well than macrophages (Figure 1L, 4B, E and S5B, C). Neutrophils are considered to be rapid responders in the first line of defense against pathogens and classically are not categorized as professional antigen presenting cells as compared to DCs, B-cells, monocytes and macrophages which have superior ability to prime naïve T cells in some settings (Lin and Loré, 2017; Reis E Sousa, 2006). However, 13% of neutrophils were MHC-II-positive (Figure 4F) suggesting they have undergone strong phenotypic maturation (Garg et al., 2017). MHC-II-positive neutrophils have recently been linked to immunogenic cell death (ICD) during which they phagocytose dying tumor cells and mediate respiratory-burst-dependent cytotoxicity against residual cells (Garg et al., 2017).

Interestingly, panobinostat was the only one of the seven drugs tested that induced substantial cell kill near the drug delivery site as indicated by significantly increased CC3 expression (Figure 4C, D). Based on our observation of a significant enrichment of MHC-II+ antigen presenting neutrophils at the panobinostat well, we hypothesized that panobinostat-mediated cell death would be immunogenic and the efficacy of this targeted therapy would be enhanced by PD-1 blockade.

We confirmed that panobinostat-induced tumor killing was immunogenic by performing a whole animal vaccination study (Ma et al., 2013; Md Sakib Hossain et al., 2018). Specifically, aliquots of E0771 and EMT6 tumor cells treated with panobinostat *in vitro* or killed by freeze thawing (negative control for non-immunogenic cell death) were injected subcutaneously into syngeneic immunocompetent mice, and then mice were then re-challenged with live tumor cells of the same type after seven days. We observed significantly increased tumor-free survival in mice immunized with panobinostat-treated tumor cells as compared to mice inoculated with untreated freeze-thawed dead cells (P value <.0001 and 0.0027 for E0771 and EMT6, respectively; Figure 4G). Consistent with this, systemic treatment of mice with panobinostat significantly increased the proportion of intratumoral CD8+ T cells as compared to stromal parenchyma (Figure 4H and Figure S5D).

Next, we tested the utility of combining panobinostat with an anti-PD-1 antibody by systemically treating mice carrying syngeneic mammary tumors with these drugs administered singly and in combination. Panobinostat alone significantly decreased the rate of tumor growth in early-stage tumors (Figure S5E) but later stage tumors did not respond to the single agent (Figure 4I, S5F). However, combining panobinostat with anti-PD-1 immunotherapy, as suggested by our MIMA studies, significantly decreased tumor growth rate and increased survival in both EMT6 and E0771 models (Figure 4I, 6H) thus indicating effective induction of antitumor immunity.

### Biomarkers of response and mechanisms of resistance help to identify early intratumor signatures of panobinostat-induced anti-tumor immunity in mammary carcinoma

Because our local and systemic studies indicated panobinostat to be increasing immunogenicity of mammary carcinoma (Figure 4, S5), we further evaluated potential mechanisms of response and resistance associated with this targeted therapy. We extended the mIHC readout to enable assessment of functional biomarkers related to induced anti-tumor immunity by adding antibodies to probe expression of calreticulin which facilitates folding of the MHC-I complex in the endoplasmic reticulum (Raghavan et al., 2013) and dictates immunogenicity of cell death (Obeid et al., 2007), intercellular adhesion molecule 1 (ICAM1) and the *β*-galactoside-binding lectin, galectin-3; which play roles in recruitment of neutrophils to tissues (Gittens et al., 2018; Yang et al., 2005) and licensing of myeloid cells (Radsak et al., 2000; Vonderheide, 2018), myeloperoxidase (MPO) to identify cytotoxic neutrophils (Patnaik et al., 2017), and MHC-I and neuropilin-1 to report on proficient antigen presentation capacity (Chawla et al., 2016; Kerros et al., 2017; Luo et al., 2018). These early in situ markers as well as PD-L1 have been directly or indirectly associated with immunogenic cell death, increased tumor CD8+ T cell infiltrate and/or immune checkpoint blockade efficacy (Aguilera et al., 2016; Guerriero et al., 2017; Hu and McArthur, 2018; Luo et al., 2018; Obeid et al., 2007; Patnaik et al., 2017). We measured the expression of these proteins at the panobinostat reservoir and refer to them in ensemble as “biomarkers of immunogenic cell death” (Figure 1D, in yellow). We also added antibodies to probe resistance mechanisms associated with breast cancer progression; specifically, the cancer stem cell (CSC) marker, Sox9 (Guo et al., 2012); the immune suppression marker arginase-1 (Geiger et al., 2016; Rodriguez et al., 2017); Keratin-14 as a marker of the tumor invasive front (Cheung et al., 2016, 2013); and matrix metalloproteinases (MMPs) and collagens as markers of extracellular matrix (ECM) processing and deposition (Sahai et al., 2020) (Figure 1D, in blue). We assessed the locations of the 17 standard cell types and the expression of the various biomarkers of response and resistance thereon and found that they were organized in distinct layers (zones) at increasing distances from the panobinostat delivery well. These zones were designated as *immediate, proximal, border, distal* and *remote* as illustrated in Figure 5B.

**Figure 5.**
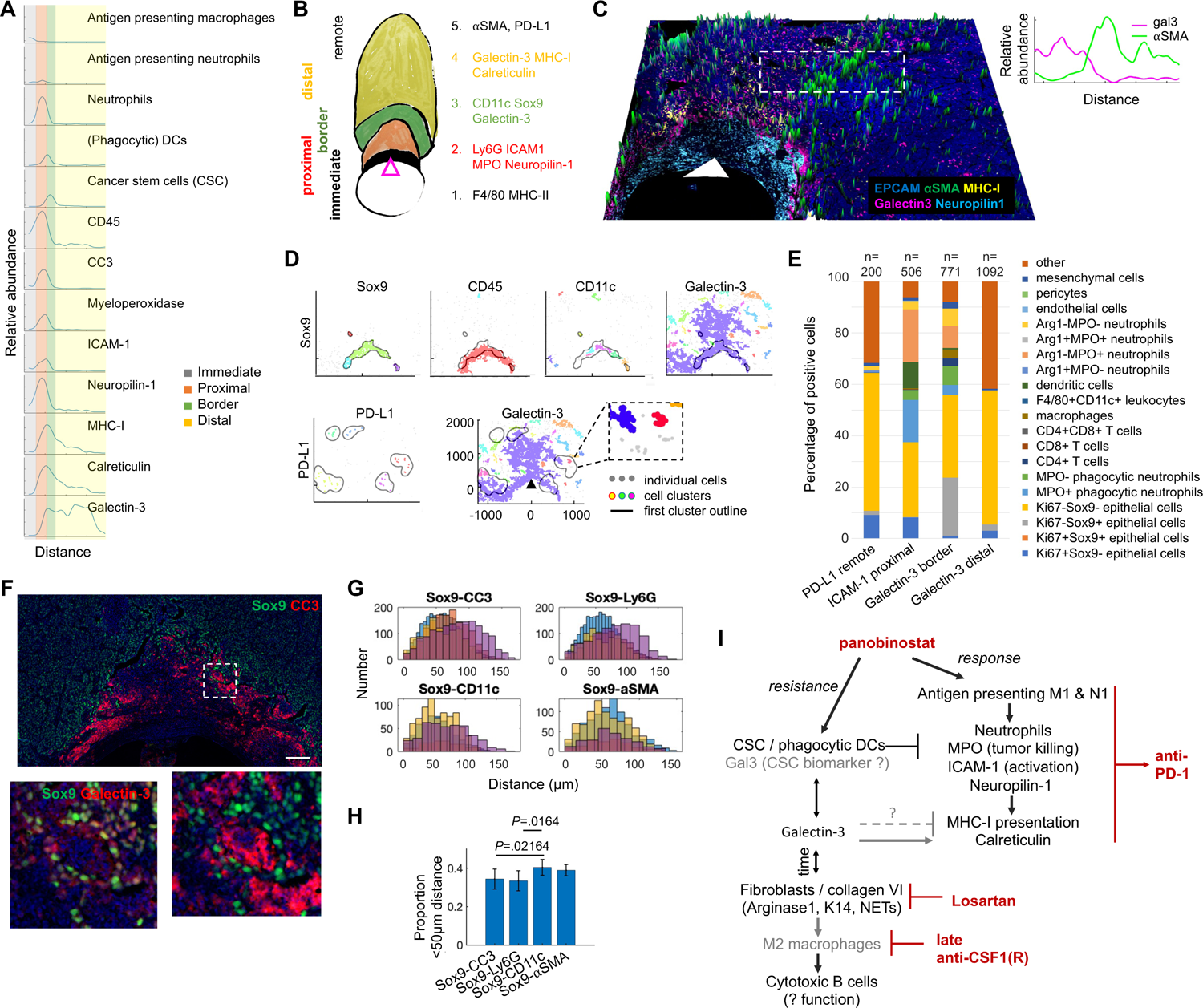
Spatial cell analyses of immunogenic cell death biomarkers and associated resistance mechanisms. (A) Profile plot of the relative abundance of standard cell types and individual biomarkers with distance from the well and overlay with the assay zones (colored vertical lines). (B) A schematic presentation of cell phenotype separation into zones with distance from the panobinostat reservoir. (C) 3D composite image showing biomarkers associated with immunogenic cell death and *α*SMA barrier limiting the propagation of these biomarkers presented in green. The white arrow indicates the source and direction of the panobinostat release. A profile plot of relative cell abundance at the depicted area (dashed) is presented top right. (D) Distance-based clustering of Sox9, CD45, CD11c and galectin-3 positive cells (top) and PD-L1 and galectin-3 positive cells (bottom) in XY coordinates with overlay (black line) on Sox9 and PD-L1 cluster border, respectively. Individual clusters were identified by a minimum 10 cells within a maximum 50μm distance for all but PD-L1 marker which clustered with a maximum distance set to 150μm. (E) Percentages of cells expressing biomarkers of ICD on standard cell types presented in form of a stack bar graph. The number of cells quantified is presented above the bar.(F) A composite image showing mutually exclusive staining of Sox9 and CC3; and co-expression of Sox9 with galectin-3 (bottom left image). Scale bar; 100μm. (G, H) Number of Sox9+ pairwise distances with other marker positive cells presented in form of a histogram (G) and bar graph showing average proportion of Sox9 pairwise distances which were less than 50µm (H). n=4 ROIs of 175µm diameter in the border assay zone. Significance was determined by paired two tailed t test. (I) Line diagram of a proposed panobinostat mechanism of action determined by MIMA and sites of intervention (depicted in red). Phenotypes that remain to be tested/validated are presented in gray color. All experiments were performed using the MMTV-PyMT mice with late stage spontaneously growing tumors and a three-day device implant.

Antigen presenting macrophages were located in the *immediate* region at the panobinostat well such as described above (Figure 5A, B). The *proximal* region was populated predominantly by neutrophils. These and other cells expressed MPO, ICAM1 and neuropilin-1 (Figure 5A-C, Figure S5C, SA, B). More than half (65%) of Ly6G+ neutrophils were positive for MPO (Figure S6C) suggesting they have cytotoxic capacity. Co-treatment with panobinostat and an anti-Ly6G antibody decreased panobinostat mediated cell death implying that these neutrophils have anti-tumor function as a result of the drug’s mechanism of action (Figure S6D). These results indicate that recruitment/induction of cytotoxic neutrophils is an important mechanism by which panobinostat causes cell death. We expected ICAM-1 to be expressed in the perivascular space (Patnaik et al., 2017). However, it was mostly expressed on neutrophils (44%), tumor cells (37%) and DCs (10%, Figure 5E). The expression of ICAM-1 by myeloid cells including neutrophils suggests that they might be activated and capable of priming T cells to induce anti-tumor immunity (Banchereau and Steinman, 1998; Radsak et al., 2000). Neuropilin-1 is a molecule with pleiotropic function and is mostly pro-tumorigenic in other cancers (Graziani and Lacal, 2015; Matkar et al., 2016; Overacre-Delgoffe et al., 2017); however, in breast cancer, it was recently reported to improve class I antigen presentation machinery and cross-presentation (Chawla et al., 2016; Kerros et al., 2017). The majority (up to 88%) of neuropilin-1 positive cells proximal to the panobinostat well were cytotoxic neutrophils (Figure 5A, C and S5C, S6E, F).

The high phagocytic and tumor-killing potential, high expression of ICAM-1 and mutually exclusive expression of the immune suppressive molecule arginase-1 on the panobinostat induced neutrophil population (Figure S5C, S6A) indicate these are likely anti-tumor (reported also as N1) rather than protumor (N2) neutrophils (Fridlender et al., 2009; Shaul et al., 2016). These results raise the possibility that neutrophils induce increased MHC-I expression on neoplastic cells and suggest that neuropilin-1 may be a novel biomarker of anti-tumor neutrophils in BC – hypotheses that remain to be functionally tested.

The *distal* region was populated predominantly by tumor cells expressing high levels of galectin-3, MHC-I, calreticulin and PD-L1 (Figure 5A-E; S5C and S6); the latter being expressed >500um from the well at the outer border of a galectin-3 rich region (Figure 5D, S6A). Relative increase of MHC-I and calreticulin expression on the surface of cells was present in a gradient pattern that was highest in proximal cell death/neutrophil regions and decreased with distance from the panobinostat well (Figure 5A and S6A). We observed a similar spatial cell organization in a different genetically engineered BC mouse model – ErbB2*Δ*Ex16 (Turpin et al., 2016) (Figure S7A, B) highlighting the generality of this phenomenon.

Two apparent cellular barrier transition zones co-evolved with the biomarkers of immunogenic cell death: (i) first at the outer *border* of the proximal neutrophil rich region composed of cancer stem cells associated with DCs limiting propagation of cell death; (ii) second at the outer border of galectin-3, MHC-I and calreticulin-rich region *remotely* from the well (Figure 5A-C).

The *border* region was populated by CSCs defined as Epcam+CD45-PyMT+ cells with nuclear expression of Sox9 (Guo et al., 2012) which were intermixed with CC3+ dying cells and cell debris (Figure 1L, S5B, C). CSCs have been reported to have self-renewal and tumor-initiating capacity and often exhibit resistance to therapy (Jeselsohn et al., 2017; Xue et al., 2019).

Importantly, cellular expression of CC3 and Sox9 staining was mutually exclusive (Figure 5F and Figure S5C) providing direct *in vivo* evidence that the CSCs were resistant to the most potent tumor killing therapy in our screen. Inversely, galectin-3 and Sox9 were co-expressed in many areas in the border region (Figure 5F) and we found that 22% of galectin-3+ cells were CSCs (Figure 5E). This indicates galectin-3 might be another biomarker enriching CSCs in breast cancer. Finally, macroscopic profile plots of relative cell abundance (Figure 5A), distance-based cluster analyses (Figure 5D) and pairwise proximity measurements in Sox9 microcultures (Figure 5G, H and S6G, H) showed that, among immune cells, CD11c+ dendritic cells were preferentially located in close proximity to the CSCs suggesting functional interactions between the two cell types.

Spindle-shaped *α*SMA+ cells – likely activated fibroblasts – consistently appeared in the *remote* region (Figure 1L, 5C, S6F). We speculate that these cells may act as a barrier to physically restrict cellular and molecular movements since they form a sharp boundary that appears to limit the galectin-3 signal propagation (Figure 5C, top right). This *α*SMA barrier became increasingly prominent at day 8 and more phenotypes emerged at that time that have been associated with mechanisms of resistance; including K14+ cells comprising a tumor invasive front, immune suppressive expression via arginase-1, strong deposition of Collagen VI, Ki67+Ly6G+ (likely neutrophil extracellular trap (NET) formation (Albrengues et al., 2018)), and accumulation of Sox9 cancer stem cells in close proximity to both dendritic cells and fibroblasts (Figure S7C).

The transition of galectin-3 from diffuse spreading across the distal zone (Figure 5C, D, S5C and S6A, F, G), into a sharp barrier (Figure S7C) suggests it may be a resistance marker and critical component spatially connecting fibroblasts to the CSCs/DCs niche.

Pro-tumorigenic macrophages with low expression of CSF1R (Figure S7C) and sparse single cytotoxic B cells expressing CD45R, granzyme B, galectin-3 and calreticulin (Figure S7D) appeared at the outer edge of the resistant zone remotely from the well at day 8 of panobinostat exposure. Expression and spatial association of galectin-3 with functionally distinct – both response and resistance – phenotypes (calreticulin, antigen presentation, cytotoxicity, PD-L1, stem cells, immune suppression) suggest a broad and possibly time-dependent function of this protein indicating that targeting galectin-3 during immunogenic cell death should be carefully considered (Figure S6D). While quantitative evaluation of the spatial cell composition of immunogenic cell death across multiple time points is outside the scope of this study, the appearance of spatially segregated phenotypes after three and eight days shows that spatial assessment of cellular responses with increasing distance from the reservoir provides insight into the sequence of emerging cellular events that follow treatment (Figure S7A) and ultimately may be reverse engineered to devise effective treatment schedules (Figure 5I, Figure S8). The cause-consequence spatial cell associations describing the early in situ events of induced anti-tumor immunity in BC are summarized in Figure 5I.

### Combination of panobinostat, venetoclax and anti-CD40 immunotherapy maximizes tumor killing and immune surveillance in mammary carcinoma

We used information from additional MIMA analyses to identify drugs that would enhance panobinostat mediated tumor killing and immune-modulation. In one analysis, we assessed effects by combining panobinostat with other drugs in individual MIMA reservoirs. We decreased the baseline concentration of panobinostat to approximately 25% of the original concentration in order to facilitate identification of highly synergistic combinations. We then measured the relative increase of cell death (CC3) and leukocyte density (CD45) compared to control and panobinostat treatments after 3 days. Combinations of panobinostat with venetoclax, doxorubicin and palbociclib were highly effective at increasing cell death and leukocyte density. Combining panobinostat with venetoclax significantly potentiated immune modulation, while its combination with doxorubicin significantly increased tumor kill (Figure 6A). In a second analysis, we delivered single drugs in adjacent wells and assessed the effects in regions of drug overlap between the wells (Figure 6B) over the course of eight days. These studies were carried out in orthotopic MMTV-PyMT derived mammary tumors. Orthotopic models exhibit a lower level of non-immune stroma at baseline (Dunn et al., 2004; Yang et al., 2017) and the eight day exposure was chosen to provide sufficient time for the drugs to diffuse to create a zone of drug overlap. We focused on the combinations of panobinostat with venetoclax or palbociclib in subsequent experiments as panobinostat/doxorubicin combination treatment efficacy had been evaluated previously (Budman et al., 2012).

**Figure 6.**
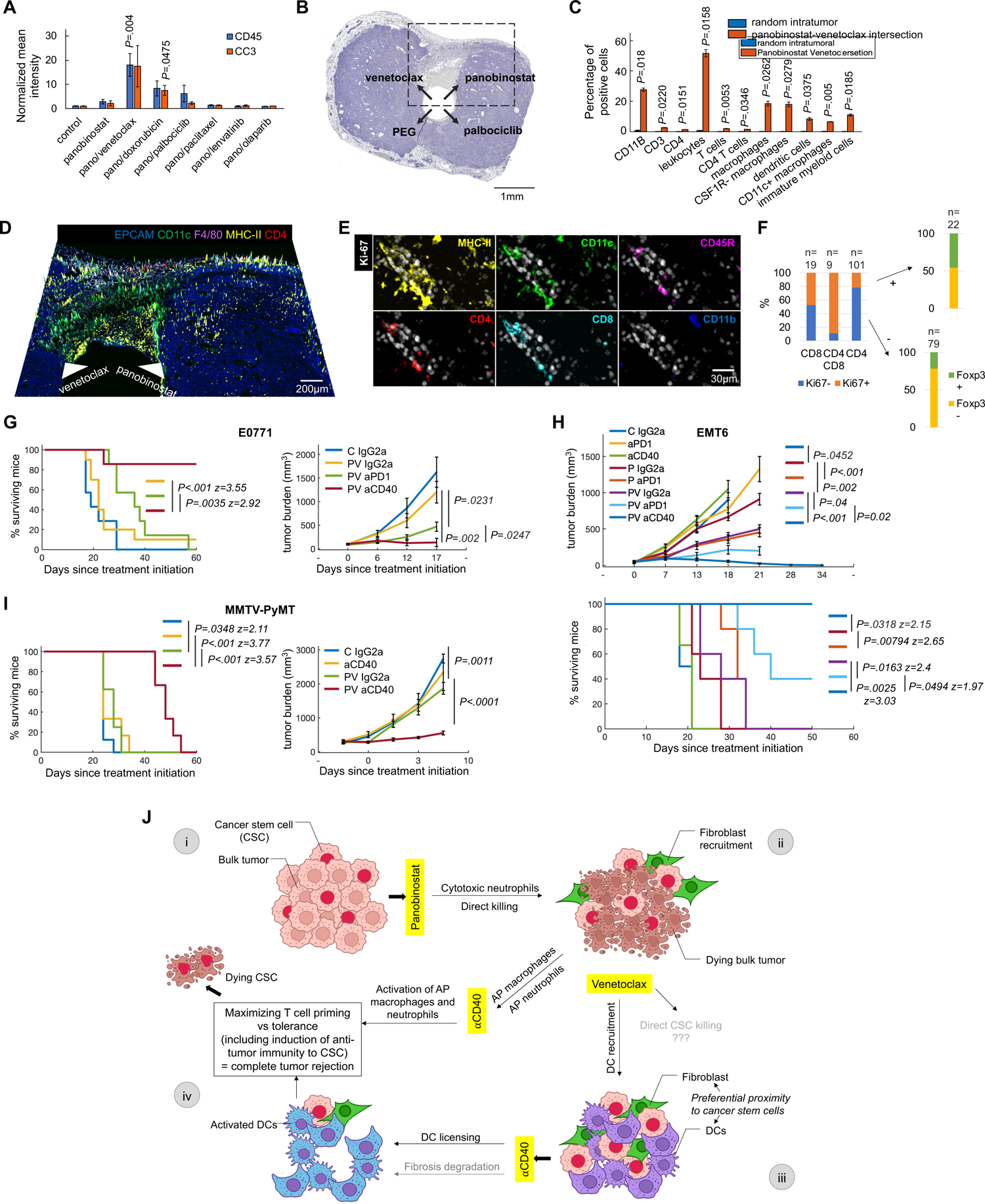
Efficacy of the triple combination of panobinostat, venetoclax and anti-CD40 immunotherapy in mammary carcinoma and rationale for the combination. (A) Effects of drug combinations on CC3 and CD45 expression as measured by IHC at three days of exposure. Graph shows mean ± s.e normalized signal intensity. n=3-10 wells from at least 3 tumors from 2-3 MMTV-PyMT mice with spontaneous tumors per treatment; significance was calculated by independent two-sample t-test with equal variance. (B) Macroscopic view of a hematoxylin-stained tumor tissue section showing the intersection between two drugs. Drug release sites are shown by black arrows. The device was implanted for eight days in MMTV-PyMT tumor induced by orthotopic implant into mammary fat pad of syngeneic mice. Note the tumor cleared region lacking nucleated cells at the intersection of panobinostat and venetoclax. (C) Quantification of single cell events using individual markers and marker combinations for standard cell types. Bars are mean ± s.e.; n=2 reservoirs at the intersection of panobinsotat and venetoclax. Significance was calculated by paired sample one tailed t-test. Only significantly enriched cells are presented. For quantification of all cells, see Figure S9C. (D) A five-color 3D composite image showing key response markers induced at the intersection of panobinostat and venetoclax. White arrows indicate the source and direction of the drug release. CD11c dendritic cell marker is presented in high view. Scale bar; shown. (E, F) Limited infiltration of CD4+, CD8+ T cells and CD45R+ B cells localized to MHC-II antigen presenting CD11c+ dendritic cells lacking CD11b pan-myeloid marker expression (E) and quantification of their Ki67 proliferative and Foxp3 regulatory potential (F). (G, H, I) Survival rate (left and bottom; 100% to 0%) and tumor burden measurements (right and top) over time in E0771 (G), EMT6 (H) orthotopically induced tumor bearing mice and MMTV-PyMT mice with spontaneously growing tumors (I). C, control; P, panobinostat, PV, panobinostat-venetoclax combination. Treatment schedules and doses match those in Figure 3F and 4H except the doses for panobinostat and venetoclax were decreased to 11.5mg/kg and 18mg/kg, respectively, when drugs were combined. For survival rate, P-value was calculated by log-rank (Mantel-Cox). For tumor burden, line graphs are mean ± s.e. per timepoint; n= 8-12 mice, and 6-12 tumors and 6-8 mice per group in (G), (H) and (I), respectively. Significance was calculated by unpaired two-tailed t-test with equal variance. (J) Hypothetical model of response for panobinostat/venetoclax/anti-CD40 triple combination treatment efficacy in breast cancers. Briefly, the tumor is composed of bulk tumor and cancer stem cells (i). Panobinostat induces immunogenic cell death of the bulk tumor while cancer stem cells remain resistant in the tumor microenvironment (ii). Venetoclax induces recruitment of dendritic cells in close proximity to cancer stem cells (iii). We hypothesize that if CD40 ligation induces licensing of dendritic cells which captured and processed antigen from neighboring CSCs, the triple combination potentiates CSC-specific anti-tumor immunity leading to complete tumor rejection (iv).

Remarkably, the combination of panobinostat and venetoclax resulted in complete clearance of tumor cells at the intersection of the two drugs (Figure 6B, Figure S9A, B). The observed TME responses included significant enrichment of macrophages, dendritic cells, immature myeloid cells and CD4+ T cells (Figure 1H, 6C, D and Figure S9C). A cluster of MHC-II positive DCs co-expressing Ki67+ CD4 and CD8 appeared near stroma and assay region (Figure 6E and S9A) suggesting the drug combination induced intra-tumoral T cell infiltration and stimulation (Broz et al., 2014). However, this event was rare, and the vast majority of the panobinostat/venetoclax assay area was dominated by myeloid cells (Figure 1H, 6C and S9C) implying that a myeloid targeting agent rather than ICB could optimally exploit the panobinostat/venetoclax induced tumor microenvironmental state.

Since CSF1R positive protumorigenic macrophages were not significantly enriched at the intersection of wells (Figure S9A-C) and the anti-CD40 agonist antibody can positively modulate DCs, immature myeloid cells (Hegde et al., 2020; Sotomayor et al., 1999) as well as resting and proinflammatory macrophages (Verreck et al., 2006), we tested the possibility that anti-CD40 immunotherapy would further increase the efficacy of the panobinostat/venetoclax combination by licensing the accumulated myeloid cell subsets as described above. We tested this by systemically treating mice bearing EMT6 and E0771 orthotopic tumors and compared the responses to those obtained using a panobinostat/venetoclax/anti-PD-1 combination. Treatment with panobinostat/venetoclax/anti-PD-1 significantly reduced the tumor burden as compared to dual panobinostat/venetoclax and panobinostat/anti-PD-1 (Figure 6G, H) treatments with survival rates of 40% in mice bearing EMT6 tumors (Figure 6H). The triple combination of panobinostat/venetoclax/anti-CD40, however, was superior and eliminated visible tumors in 100% of EMT6 tumors and 85% of E0771 tumors, respectively (Figure 6G, H). The antitumor effect of panobinostat/venetoclax/anti-CD40 against spontaneous tumors arising in the MMTV-PyMT model, inhibited tumor progression and doubled the overall survival (Figure 6I).

We note that none of the combination treatments in whole animal studies were associated with adverse events, likely because we used lower concentrations of drugs than published previously (Table S5). We measured underconditioned body score (level 2; as established by (Morton and Griffiths, 1985)) associated with palbociclib monotherapy treatment; and two out of eight mice died in the anti-CD40 monotherapy group. Lethal toxicity of anti-CD40 used as a single agent was previously reported due to a shock-like syndrome (Van Mierlo et al., 2002) and our data also suggest this immunotherapy is tolerable only with prior administration of anti-cancer agent(s).

Overall, these results suggest the triple combination of panobinostat, venetoclax and anti-CD40 immunotherapy as a highly synergistic therapeutic strategy for long term breast cancer control.

## DISCUSSION

Much research is now underway to develop synergistic multi-drug cancer treatment strategies that directly target neoplastic cells, enhance anti-tumor immune activation, normalize tumor vasculature and/or favorably alter aspects of the tumor microenvironment to improve tumor control. Successful strategies increasingly use combinations of drugs, each of which impacts one or more components of the TME. We developed the MIMA system to efficiently develop effective combination regimens by assessing tumor cell control and drug-induced changes in immune and stromal composition, architecture and function that correlate with overall antitumor efficacy. MIMA attributes include an IMD for delivery of nanoliter doses of multiple drugs, each delivered into a spatially separate region of a single living tumor growing in an immunologically intact host; a 30+ multiplex IHC analysis platform that provides a comprehensive description of tumor and stromal responses to each localized drug treatment and the functional status of selected immune and other tumor and stromal cells. The approach provides a highly precise, multiplexed platform to systematically identify candidate biomarkers of response and quantify cell interactions and to inform on treatment sequencing (Figure S8).

While majority of our studies were performed using a three day long microdevice implantation, the in-dwelling time of the IMD can be varied from a few hours to many days in order to capture a range of tissue responses and TME reorganization. Importantly, the approach is minimally toxic even when testing multiple therapies in a single tumor. The local drug doses match concentrations that are achieved during systemic treatments but are systemically insignificant such that drug toxicity is negligible. The focal drug delivery begins at the time of implantation, and follows a characteristic diffusion gradient controlled by the PEG polymer formulation in a more-or-less radial direction away from each drug delivery well (Jonas et al., 2015). The focal delivery can then be treated as a spatial and temporal perturbation. Analyses of the responses produced by devices left in place across a range of time points provide data about drug induced changes in cellular densities, molecular phenotypes, cell motility and functional interactions. In addition, analyses of the regions between drug delivery wells where drugs are allowed to overlap via diffusion serves as a measurement of the effect of the combination of those drugs. Since distances from the drug delivery wells reflect recruitment events, computational modeling of these patterns can provide actionable information to guide the development of effective drug doses and schedules.

Although not pursued explicitly in this study, the timing of combination immunotherapies in whole animals was estimated based on immune component responses at increasing distances from the drug delivery wells (Figure S8). Many of these effects are difficult or impossible to study in animal models treated systemically, due to heterogeneous and indeterminate drug distribution that can vary greatly over different regions of a tumor. Importantly, we validate the significance of the locally observed response phenotypes from MIMA studies by reverse-engineering combination treatments involving targeted and immuno-therapies (Table S4) that demonstrate synergistic anti-tumor efficacy in systemic studies. This is particularly important since it enables systems level studies of the effects of multiple drugs in a single organism, and leads to accurate predictions of responses to systemic treatments in animal models with intact immune systems. Furthermore, recent work by Jonas et al has demonstrated that IMD applications are safe and feasible in patients across multiple cancer indications including breast cancer, prostate cancer, T cell lymphoma and glioblastoma (Dominas et al., 2021). It may become feasible then to use the MIMA assay in patients to measure a range of combination regimens in each patient to guide rational treatment design on a personalized basis.

Although intended as proof-of-concept that local nanodose drug phenotypes can effectively guide systemic treatment strategies, we have already identified specific therapeutic strategies that warrant clinical consideration. One finding is that the CDK4/6 inhibitor, palbociclib, recruits a significant number of CSF1R+, MHC II-protumorigenic macrophages that appear to induce formation of CD31+ endothelial/pericyte networks that contribute to neovascularization and provide nutrients to support tumor cell proliferation. These results provide direct evidence of how specific changes in tumor microenvironmental states induced by monotherapy can mediate acquired resistance. We hypothesized that this protumorigenic resistance phenomenon may be overcome by combining palbociclib with anti-CSF1R antibody. Our test of this concept in the EMT6 model demonstrated that systemic treatment with this drug combination significantly reduced tumor growth.

Our studies of venetoclax demonstrated that this BCL-2 inhibitor induced formation of phenotypically distinct clusters of CD11c+ dendritic cells associated with immature myeloid cell and endothelial cell enrichment. Many of the dendritic cells were Epcam+, CD45-suggesting that they were phagocytic, while others shared morphological features of unstimulated myeloid cells. However, only a small fraction of these cells were MHC-II positive and thus were likely limited in their ability to respond to available tumor antigens. This finding led us to add an anti-CD40 immunotherapy to increase antigen presentation, maturation and activation (aka licensing) in a population that was already poised to have antitumor activity. Our test of this hypothesis in the E0771 model showed that systemic treatment with a combination of venetoclax and an anti-CD40 agonist indeed reduced tumor growth rate and increased overall survival as predicted (Figure 3G).

Our demonstration that the pan HDAC inhibitor, panobinostat, increased immunogenicity (by induction of immunogenic cell death and antigenicity; (Yatim et al., 2017)) of mammary carcinomas when administered locally or systemically in four different animal models of breast cancer corroborates previous studies showing the importance of HDAC inhibitors as pleiotropic effectors of many immune surveillance processes (Conte et al., 2018). This immune component is important in breast cancer since many patients do not benefit from treatments including immune checkpoint blockade (Force et al., 2019). However, the exact mechanisms by which HDAC inhibitors influence immune surveillance are still being elucidated (Conte et al., 2018) and likely vary between inhibitors. Treatment of EMT6 and E0771 model tumors with panobinostat plus anti-PD-1 increased survival duration and reduced tumor growth rate relative to treatment controls or to treatment with panobinostat alone (Figure 4H, 6H). However, these treatments did not achieve long term tumor control and in vaccination studies – not all mice in either EMT6 and E0771 model rejected the tumor post re-challenge (Figure 4G). These studies suggest that resistance mechanisms exist that might counter the full potential panobinostat mediated antitumor immunity and thus we explored this treatment condition in more detail.

Our studies indicate that panobinostat enhances anti-tumor immunity by recruitment of antigen presenting macrophages, cytotoxic and antigen presenting antitumor neutrophils and ICAM-1 positive myeloid cells leading to upregulation of MHC-I and calreticulin expression on tumor cells. However, this is counterbalanced by the emergence of multiple potential resistance mechanisms including preferential clustering of dendritic cells with cancer stem cells, the formation of fibroblast/ECM barriers to treatment and antigen presentation, recruitment of immune suppressive cells (arg-1), development of highly metastatic tumor cells expressing K14, NETs and CSF1R+ cells. Recruitment of CSF1R+ cells at a late time point indicatings sequential or alternating administration of HDAC inhibitors with CSF1/CSF1R targeting agents may be efficacious while simultaneous dosing is not (Guerriero et al., 2017)) (Figure 4C and S8). Our studies also identified neuropilin-1 and galectin-3 as candidate biomarkers that may inform on panobinostat mechanism of action. Neuropilin-1 appears to mark anti-tumor (N1) neutrophils that may potentiate antigen presentation in mammary carcinoma (Chawla et al., 2016; Kerros et al., 2017) while galectin-3 expression is associated with several anti-tumor endpoints including increased tumor MHC-I, calreticulin; cytotoxic granzyme B positive B cells as well as pro-tumor phenotypes (cancer stem cells, immune suppression). The validated biomarker of early induced antitumor immunity may allow early stratification of breast cancer patients to immune checkpoint blockade. This concept may be further investigated in window of opportunity treatment clinical trials.

Analyses of the tumor and its microenvironments in regions of panobinostat and venetoclax intersection revealed almost complete tumor clearance. These two targeted anticancer agents worked together to increase total intratumoral immune cell counts – panobinostat by inducing neutrophils and macrophages and venetoclax by inducing recruitment of DCs and immature myeloid cells. Importantly, all these myeloid cells can be positively modulated by anti-CD40 immunotherapy. DCs are generally thought to be the main antigen presenting cells which can activate naïve T cells to become effectors (Mempel et al., 2013). However, CD40 ligation also upregulates antigen presenting molecules and adhesion molecules such as ICAM-1 and increases the type-1 proinflammatory state to support immunogenic processes in neutrophils, resting and proinflammatory macrophages and immature myeloid cells (Oehler et al., 1998; Radsak et al., 2000; Sotomayor et al., 1999; Verreck et al., 2006) (Figure 6J). Additionally, in pancreatic cancer, this immunotherapy modulates the TME to degrade fibrosis (Long et al., 2016). It remains to be determined if such matrix remodeling of the formed fibrotic barriers could also occur in breast cancer (Figure 6J). Also, the spatial association of DCs and CSCs might be of particular importance in the panobinostat/venetoclax/anti-CD40 mechanism of action. We hypothesize that panobinostat induces anti-tumor immunity to bulk tumor while cancer stem cells remain resistant in the TME. Venetoclax induces recruitment of DCs which we have revealed to localize specifically to the CSC niche. We propose that if CD40 ligation induces licensing of DCs which already captured and processed CSC antigen, this might result in activation of CSC-specific anti-tumor immunity and eventually to complete tumor clearance (Figure 6J). Thus, we suggest that panobinostat induces antitumor immunity on the level of bulk tumor, while venetoclax/anti-CD40 induces anti-tumor immunity on the level of resistant, tumor initiating cancer stem cells. This model of response is so far hypothetical as antigen specific T cell responses remain to be critically evaluated. Nevertheless, our observations suggest that the combination of lower dose panobinostat/venetoclax/anti-CD40, and the anti-CD40 immunotherapy in general, should be considered clinically for treatment of breast cancer.

In sum, the MIMA platform described here provides a strategy to design effective combination regimens based on intratumor nanodose exposure to a range of agents, coupled with highly multiplexed phenotyping and integrated spatial analysis of tumor response to each therapy. By testing multiple therapeutic strategies in the same tumor, we can for the first time perform systems level analysis using multiple parallel pharmacological perturbations in the same organism. The low drug toxicity of intratumor nanodosing further supports clinical use of MIMA, which is currently being investigated in multiple human studies. Thus, MIMA represents a new approach to identification of effective combination regimens for individual patients on a personalized basis. Extended use of MIMA will also open new opportunities in *in silico* modeling to model dynamic drug-tumor-stromal interactions.

## ACKNOWLEDGEMENTS

We thank the OHSU Histopathology Share Resources core (Todd Camp, Joscelyn Zarceno and Cheyenne Martin) for careful FFPE processing of the MIMA tumor samples; Young Hwan Chang, Sam Sivagnanam, Eugene Manley and AeSoon Bensen for help with image processing and mIHC/cycIF staining; Tiziana Cotechini for sharing the LPA3 mouse model line; and Marilyn McWilliams and Dottie Waddell breast cancer advocates for valuable feedback during project reports. This work was supported by the NIH/NCI Cancer Systems Biology Consortium Center U54CA209988 (JWG), Susan G. Komen grant SAC190012 (JWG) and the NIH grant R01CA223150 (OJ, LMC).

## AUTHOR CONTRIBUTIONS

Conceptualization, ZT, OJ, JWG; Methodology, ZT, OJ, JWG; Software, ZT, DCB; Investigation, ZT, DCB; Data Analysis and Interpretation, ZT, OJ, JWG; Writing – original and final draft, ZT, OJ, JWG; Writing – review & editing, JEK, LMH, JLM, PJS, SWA, GBM, LMC; Resources, JLM, LMC, OJ, JWG; Funding, OJ, JWG; Supervision, OJ, JWG.

## STAR METHODS

### CONTACT FOR REAGENT AND RESOURCE SHARING

Further information and requests for resources and reagents should be directed to and will be fulfilled by the lead contact Joe W. Gray.

### EXPERIMENTAL MODELS AND SUBJECT DETAILS

#### Murine Models

Mice were purchased from The Jackson Laboratory. All animal studies were conducted in accordance with protocols approved by Institutional Animal Care and Use Committee (IACUC) at OHSU (protocol number: IP00000956). All mice were bred and housed under specific pathogen free conditions under a standard 12h light / 12h dark cycle. C57LB/6, BALB/c, and FVB/N mice were purchased from the Jackson Laboratory. MMTV-PyMT were from Dr. Lisa Coussens and purchased from the Jackson Laboratory. Virgin female mice of 8-24 weeks of age were used for all experiments.

#### Cell lines

EMT6 (mouse breast cancer) cells were purchased from American Type Culture Collection and were maintained in Waymouth’s medium with 10% FBS, and 2mM L-glutamine. E0771 (mouse breast cancer) cells were purchased from CH3 BioSystems® and were cultured in RPMI-1640 with 10% FBS and 10mM HEPES. Both cell lines were pathogen tested and were grown at 5% CO_2_ and 37C.

## METHOD DETAILS

### Experimental design

The objective of the studies in figures is to show how intact tumor microenvironment responds to local stimulus of drug release and to test whether this response was significantly different from the baseline tumor microenvironmental state in tumor region distant from the drug site. The number of independent biological replicates of each experiment (n) performed are given in the figure legends. Spatial systems analyses were designed to quantitatively define directional spatial cell dependencies and cause consequence association with distance from the reservoir translating to models of drug response. Within these models we aimed to identify therapeutic vulnerabilities to predict rational immune or TME modulating treatment combinations and their optimal schedule/sequencing.

### Microdevice implantation studies and sample collection

Nanodose drug delivery devices were manufactured and implanted as described previously in (Jonas et al., 2015). Briefly, cylindrical microdevices 5.5mm in length and 750μm in diameter were manufactured from medical-grade Delrin acetyl resin blocks (DuPont) by micromachining (CNC Micromachining Center) with 18 reservoirs 200μm (diameter) x 250μm (depth) on the outer surface. Reservoirs were packed with drugs mixed with Polyethylene glycol (PEG, MW 1450, Polysciences) polymer at the concentrations indicated in Table S1. Recommended systemic dose in cancer patients was derived from the https://rxlist.com web page to June 2017. Systemic doses ranging between 0-1mg/kg, 1-2mg/kg, 2-4mg/kg, >4mg/kg translate to 20%, 25%, 30% and 40% of drug concentration in PEG, respectively, when released from the nanowell. The calibration was determined previously using mass spectrometry measurements (Jonas et al., 2015). Pure PEG was used in control conditions. Implanting multiple devices per tumor and/or multifocal animal model can increase the throughput up to 50-70 times as compared to conventional systemic treatment studies. When two drugs were loaded into one reservoir, they were at approximately 1:1 ratio. The combination partner was loaded on the bottom of the well; panobinostat was released first. Microdevices were implanted for three days in MMTV-PyMT and ErbB2*Δ*Ex16 mice with late stage spontaneously growing tumors in all experiments but those presented in Figure 6 and S9. Tumor size was between 1.2 - 1.5cm in the longest dimension at the time of implant. Tumors were excised at three days after device implantation unless otherwise stated, fixed for 48h in 10% formalin or 4% paraformaldehyde, then perfused with paraffin. Specimen were sectioned using a standard microtome and 5μm tissue sections were collected from each reservoir. Dry FFPE tissues were baked in a 65°C oven for 30mins. Following deparaffinization with xylene and rehydration in serially graded alcohol to distilled water, slides were subjected to endogenous peroxidase blocking in fresh 3% H_2_O_2_ for 10 minutes at RT. Sections were then stained by multiplex immunohistochemistry and/or cyclic immunofluorescence (see also Figure S1B, C).

### Cyclic Immunofluorescence

Before iterative cycles of (i) staining, (ii) whole slide scanning and (iii) fluorophore bleaching, the slides were subjected to heat-mediated antigen retrieval immersed in citrate buffer (pH 5.5, HK0809K, BioGenex Laboratories Citra Plus Antigen Retrieval), then in Tris/EDTA buffer (pH 9.0, S2368, Dako Target Retrieval Solution) using Cuisinart Electric Pressure Cooker (CPC-600N1) for total of 35 to 40 minutes. Protein blocking was performed for 30 minutes RT with 10% normal goat serum (S-1000, Vector Lab) and 1% bovine serum albumin (BP1600-100) in 1xPBS. (i) Slides were incubated with primary antibody (concentrations defined in Table S2) for 2 hours at RT while being protected from light in a dark humid chamber. All washing steps were performed for 3 x 2-5 minutes in 1xPBS while agitating. Slides were mounted with SlowFade Gold antifade mountant with DAPI (S36938) using a Corning Cover Glass (2980-245). (ii) Images were acquired using Zeiss Axio Scan.Z1 Digital Slide Scanner (Carl Zeiss Microscopy) at 20x magnification after which the coverslips were gently removed in 1xPBS while agitating. (iii) Fluorophores were chemically inactivated using a 3% H_2_O_2_ and 20mM NaOH in 1xPBS for 30 minutes at RT while being continuously illuminated. The fluorophore inactivation was repeated twice with a short, 10-minute, 1xPBS wash in between. Efficacy of bleaching was imaged before antibody incubation (baseline autofluorescence) and every third to fourth cycle in average. After protein blocking, samples were subjected to the next round of staining. Single cell feature extraction was not applied to evaluate sections stained by cyclic immunofluorescence.

### Multiplex Immunohistochemistry

Before iterative cycles of (i) staining, (ii) whole slide scanning and (iii) and heat and chemical stripping of antibodies and chromogen, the slides were subjected to staining with F4/80 and CSF1R antibodies (cycle zero, no antigen retrieval, Table S3) and hematoxylin staining (S3301, Dako) for 1-5mins followed by whole slide scanning. Slides were then subjected to the first heat-mediated antigen retrieval in 1x pH 5.5-6 citrate buffer (Biogenex Laboratories, HK0809K) for 90 seconds in a low power microwave and 16 minutes in a steamer followed by protein blocking with 10% normal goat serum (S-1000, Vector Lab) and 1% bovine serum albumin (BP1600-100) in 1xPBS for 30 minutes RT. (i) Slides were incubated with primary antibodies (concentrations defined in Table S3) for 1 hour at RT or 16-17 hours at 4 degrees Celsius while being protected from light in a dark humid chamber. Signal was visualized with either anti-rabbit or anti-rat Histofine Simple Stain MAX PO horseradish peroxidase (HRP) conjugated polymer (Nichirei Biosciences) followed by peroxidase detection with 3-amino-9-ethylcarbazole (AEC). Two or three drops of HRP polymer were used for up to nickel-size or whole slide tissue sample, respectively. Timing of AEC development was determined by visual inspection of positive control tissue (Figure S1 D-F) for each antibody. All washing steps were performed for 3 x 5-10 minutes in 1xPBS while agitating. Slides were mounted with a filtered 1xPBS with 0.075% Tween20 (BP337100) using a Signature Series Cover Glass cover glass (Thermo Scientific, 12460S). (ii) Images were acquired using the Aperio ImageScope AT (Leica Biosystems) at 20x magnification after which the coverslips were gently removed in 1xPBS while agitating. (iii) Within one cycle, removal of AEC and HRP inactivation was accomplished by incubating the slides in 0.6% fresh H_2_O_2_ in methanol for 15 minutes; AEC removal and stripping of antibodies was accomplished by Ethanol gradient incubation and heat-mediated antigen retrieval such as described above between cycles (see also Figure S1B) (Banik et al., 2020; Tsujikawa et al., 2017). After washing and protein blocking, samples were subjected to the next round of staining.

### Image processing and feature extraction of mIHC images

The iteratively digitized images were co-registered using Matlab (The MathWorks, Inc., Natic, MA, version 2019b) utilizing the detectSURFFeatures algorithm from the Computer Vision Toolbox. The imperfectly registered images were additionally processed using the Linear Stack Alignment with SIFT plugin (Fiji) so that cell features overlap down to a single pixel level.

Hematoxylin-stained images were color deconvoluted for single cell nuclear segmentation to generate a binary mask using watershed function and standard image processing steps (noise removal, erosion, dilation; Fiji) (Schneider et al., 2012). AEC chromogenic signal was extracted using the NIH plugin RGB_to_CMYK to separate AEC signal into the yellow channel for improved sensitivity of IHC evaluation (Banik et al., 2020; Pham et al., 2007). Gray scale images of all proteins and the binary mask were imported to CellProfiler (version 3.1.8, Broad Institute) (Carpenter et al., 2006) to quantify single cell signal mean intensity as defined by mask which was scaled to a range 0-1. IdentifyPrimaryObjects module was used to identify nuclei from mask; MeasureObjectIntensity module measured mean intensity for each object for each protein. The mean signal intensity per cell output was imported to FCS Express 6 and 7 Image Cytometry Software (DeNovo Software) to perform multidimensionality reduction to classify “cell standards”. Gating strategies and hierarchical cell classification is presented in Figure 1E and Figures S2E and F. Polygonal gates moving around central vertex without changing the polygon shapes was used to obtain quantitatively reproducible multiplex data, batch to batch, independent of the condition measured. Positive control tissues were used to help to define single parameter threshold for positivity by manual gating. Total of 3000-5000 cells were analyzed for feature extraction in the assay area located above the drug releasing site with ± 300 total cells for paired, experimental vs control, region. Minimum population proportion within 5% margin of error and 95% confidence level was set to 0.75% (represents 12 cells) to discriminate noise from specific cell enrichment induced by e.g. increased protein expression or cell recruitment into the assay region. Experimental condition of the assay area was compared to random control intratumoral region located perpendicular and/or far from the drug-releasing reservoir. To obtain greater control over cofounding variables, paired sample one tailed t-tests were used to determine enrichment of induced TME states. Percentage of positivity and significance was presented in form of a heatmap or bar graphs. Quality of the single cell data was ensured by excluding deformed (folded), lost or unevenly stained tissue (border effects). The assay area was determined by the first 3000-5000 cells above the well excluding these deformed regions. Single cell data from FCS Express was extracted in data grid to Matlab for downstream spatial systems analyses. In computed images, neutrophils are presented independent of the Epcam± status.

### Spatial Systems Analyses

Distance based cluster function finds clusters in a set of spatial points expressed in XY space (adapted and modified from Yann Marcon; Matlab October 2019). The clustering is based on Euclidean distance between the points (cells). The function does not require the number of clusters to be known beforehand. Each cell clusters with the closest neighboring cell if distance between the two cells is shorter than the defined threshold. Minimal number of cells per cluster are defined by user. The function outputs non-clustering cells in gray color while each cluster meeting the defined parameters (minimal number of cells within maximum distance range) are presented in randomized colors. Clusters within the maximum defined distance merge and share one color. Number of clusters and total coverage in the assay area was calculated using distinct cluster sizes (defined by minimal number of cells within maximum distance range, Figure S3F) for control PEG and palbociclib which identified that cells cluster in response to treatment if minimum 10 cells are present within maximum distance rage 30-75μm. Cluster parametrization using as few as 5 cells and distances as large as 100μm resulted in treatment non-specific cluster formation in PEG negative control. Treatment specific cluster formation with cluster definition of minimum 10 cells within 50μm distance was generalizable to all marker and standard cell types which was confirmed in panobinostat condition by comparing assay area and distal region side by side in one field of view (Figure S6G). This treatment specific cluster parametrization was applied in downstream analytics to identify hotspots/zones of interest (e.g*. proximal*, *border*, *distal*, network adjacent, CD11c+ DC clusters) in an objective, biology driven, manner.

For the relative abundance profile plot, marker positive cells and the standard cell types were extracted to XY coordinate space, signal was blurred using Gaussian Blur filter and relative abundance of positive cells was displayed with distance from the well in a profile plot as outlines in Figures S3B and S6A. A moving average filter with 50μm; and 100μm window size (movmean function; Matlab) was additionally applied to smoothen the feature signal for palbociclib and panobinostat condition, respectively. Signal in the profile plots was not scaled.

Inside the hotspot, spatial (geographical) interactions between marker positive cells were determined by proximity measurements in local microculture by using the pdist2 function in Matlab (MathWorks, Inc., Natic, MA, version 2019b) which returns the distance of each pair of observations (positive cells) in X and Y using metric specified by Euclidean distance. Random circular regions of 175µm diameter (defined by Figure S6H) were selected in the border, cancer stem cell, zone of the panobinostat assay area and Euclidean distance was measured between Sox9+ and other marker positive cells. The number of distances was presented in form of a histogram. To quantify spatially interrelated phenomenon, proportions of distances lower than 50μm (as defined by distance-based cluster analyses) was compared between different cell pairs (e.g. Sox9+/Ly6G+ vs Sox9+/CD11c+).

Extended hierarchical cell classification was applied to characterize the significantly enriched cell phenotypes forming zones of interest which were outside the standard cell type classification (e.g less differentiated macrophages or phagocytic DCs). Probe combination, number of cells analyzed within number of clusters are defined in the figures and figure legends.

2D composite and 3D composite images were presented by using Fiji (Schneider et al., 2012) and QiTissue Quantitative Imaging System (http://www.qi-tissue.com).

The spatial systems analyses were used to identify drug models of response (presented as line diagrams) and the identified therapeutic vulnerabilities were tested in whole animal studies.

### Whole animal treatment studies

MMTV-PyMT transgenic mice that were 80 days old were randomized and included in the study when their total tumor burden was between 150-550mm^3^ (treatment initiation). For the orthotopically induced tumor models of mammary carcinoma, EMT6 (0.5 x 10^6^ in 1xPBS per site), E0771 (0.5 x 10^6^ in Corning matrigel per site) and primary tumor derived LPA3 (0.8 x 10^5^ in Corning matrigel per site) cells were injected into the #4 mammary fat pad of female virgin C57LB/6, BALB/c, and FVB/N mice, respectively. One tumor was induced in the E0771, LPA3 models and two tumors were induced in the EMT6 model. Caliper measurements were used to calculate the tumor volumes using formula length x width^2^ / 2. Treatments were initiated when total tumor burden was between 60-150mm^3^ or as defined in the figure legend (Figure S5E, F). For all models, the endpoint was determined by tumor volume above 2000mm^3^ in two consecutive measurements or one measurement above 2200mm^3^. Treatments were administered by intraperitoneal injection. Dose, schedule and duration are indicated in the respective figures and figure legends. Treatment schedule was estimated depending on the location of the targetable cell phenotype in proximity to the well or more distal from the drug source. E.g. cells in the *immediate proximity* to the drug well at 3 days of exposure were likely recruited first to the drug assay area thus early targeting (pre-treatment) of these cells is preferred. Inversely, cells located in *distal* regions at late timepoints (e.g. day 8) should be targeted by posttreatment approach. See also Figure S8. Diluent and IgG2a isotype control (BioXCell) concentrations were equivalent to the highest dose of the respective drug used in each experiment.

The mice were monitored daily to determine any possible effects on the general condition of the animals using parameters as established by (Morton and Griffiths, 1985). The guidelines for pain, discomfort and distress recognition were used to evaluate weight loss, appearance, spontaneous behavior, behavior in response to manipulation and vital signs. Specifically, general appearance (dehydration, missing anatomy, abnormal posture, swelling, tissue masses, prolapse) skin and fur appearance (discoloration, urine stain, pallor, redness, cyanosis, icterus, wound, sore, abscess, ulcer, alopecia, ruffled fur), eyes (exophthalmos, microphthalmia, ptosis, reddened eye, lacrimation, discharge, opacity), feces (discoloration, blood in the feces, softness/diarrhea), locomotor (hyperactivity, coma, ataxia, circling) were monitored to determine loss of body condition (BC) score, namely: BC 1 (emaciated) score applied when skeletal structure was extremely prominent with little or no flesh/muscle mass and vertebrae was distinctly segmented; BC 2 (under-conditioned) score applied when segmentation of vertebrate column was evident, dorsal pelvic bones were readily palpable and muscle mass was reduced; BC 3 (well-conditioned) applies when vertebrae and dorsal pelvis were not prominent/visible, and were palpable with slight pressure. Loss of BC was also considered when anorexia (lack or loss of appetite) or failure to drink; debilitating diarrhea, dehydration/reduced skin turgor; edema, sizable abdominal enlargement or ascites, progressive dermatitis, rough hair coat/unkempt appearance, hunched posture, lethargy, loss os righting reflex, neurological signs or bleeding from any orifice appeared in treated mice. Majority of treated groups were well-conditioned (BC score 3); less than 20% of mice in each group experienced mild diarrhea for up to 2 days once during the course of treatment (typically post first or second therapy administration). Mice receiving palbociclib monotherapy were under-conditioned (BC score 2) starting from day 3 till the end of the treatment. Two out of eight mice in the MMTV-PyMT model died within 1-3 days after first injection of *α*CD40 immunotherapy when administered as single agent. Surviving mice receiving Venetoclax/*α*CD40 combination experienced fur graying to different degree starting approximately four weeks post treatment. No signs of pain, discomfort or distress were observed in the surviving mice. Emaciated (BC score 1), over-conditioned (BC score 4) nor obese (BC score 5) were observed in our studies. LPA-3 mice become obese with tumor development but this sign was independent of administered treatment (treatment naïve).

To measure CD8+ T cell infiltration inside the tumor bed, ErbB2*Δ*Ex16 mice with spontaneously growing tumors were intraperitoneally injected with panobinostat (15mg/mg) on day 0, 2 and 4. Tumors were extracted at day 7, were FFPE processed and were stained for Epcam and CD8 to compare rate of intratumoral (Epcam+) vs stromal (Epcam-) CD8+ T cells in panobinostat treated vs control (diluent) treated tumors.

### Vaccination study

EMT6 and E0771 cells in tissue culture were treated with a soluble drug panobinostat at 5μM concentration when they would reach 60-70% confluency. After two days the cells (80-90% death rate) were harvested and were injected subcutaneously (total 2-3 x 10^6^ cells) into lower left flank of BALB/c and C57Bl6 mice, respectively. Cells freeze-thawed three times served as negative control for non-immunogenic form of cell death. After 7-8 days, the mice were re-inoculated by injecting living cells orthotopically into one #4 mammary fat pad (total 0.5 x 10^6^ cells) and tumor appearance was monitored by minimal tumor size approximately 5mm and 3.5mm in the longest dimension for E0771 and EMT6 model, respectively (palpable tumors). We note the E0771 tumors after re-challenge appeared at the primary subcutaneous site and no tumors were developed in the orthotopic site.

### Statistical analysis

All data are combined from two to three independent experiments, unless specifically noted. To accomplish randomization for systemic mouse experiments, animals were sorted by a blinded investigator and then groups were assigned. Each group was checked post-hoc to verify no statistical significance in average starting tumor size. There was no sample-size estimation of in standard drug treatment experiments. Data are shown as mean ± SEM, unless otherwise noted. For tumor growth rate, significance was calculated by unpaired two-tailed t-test with equal variance. For survival and tumor free analyses, Kaplan-Meier curves were generated to demonstrate time to event and log-rank (Mantel-Cox) test was used to evaluate statistical significance.

## SUPPLEMENTAL INFORMATION

### SUPPLEMENTARY TABLE LEGENDS

Table S1. Drug list and drug concentration calibration used in the MIMA system; Related to Figure 1.

Table S2. Antibody order, catalog and concentration used in mouse multiplex IHC; Related to Figure 1.

Table S3. Antibody order, catalog and concentration used in mouse cycIF; Related to Figure 1.

Table S4. Rationale to select effective immunotherapy or non-immune stroma modulating combination partner based on the TME signature induced by primary treatment.

Table S5. Comparison of systemic drug dosing in our and the previously reported studies.

### SUPPLEMENTARY FIGURE LEGENDS

**Figure S1.**
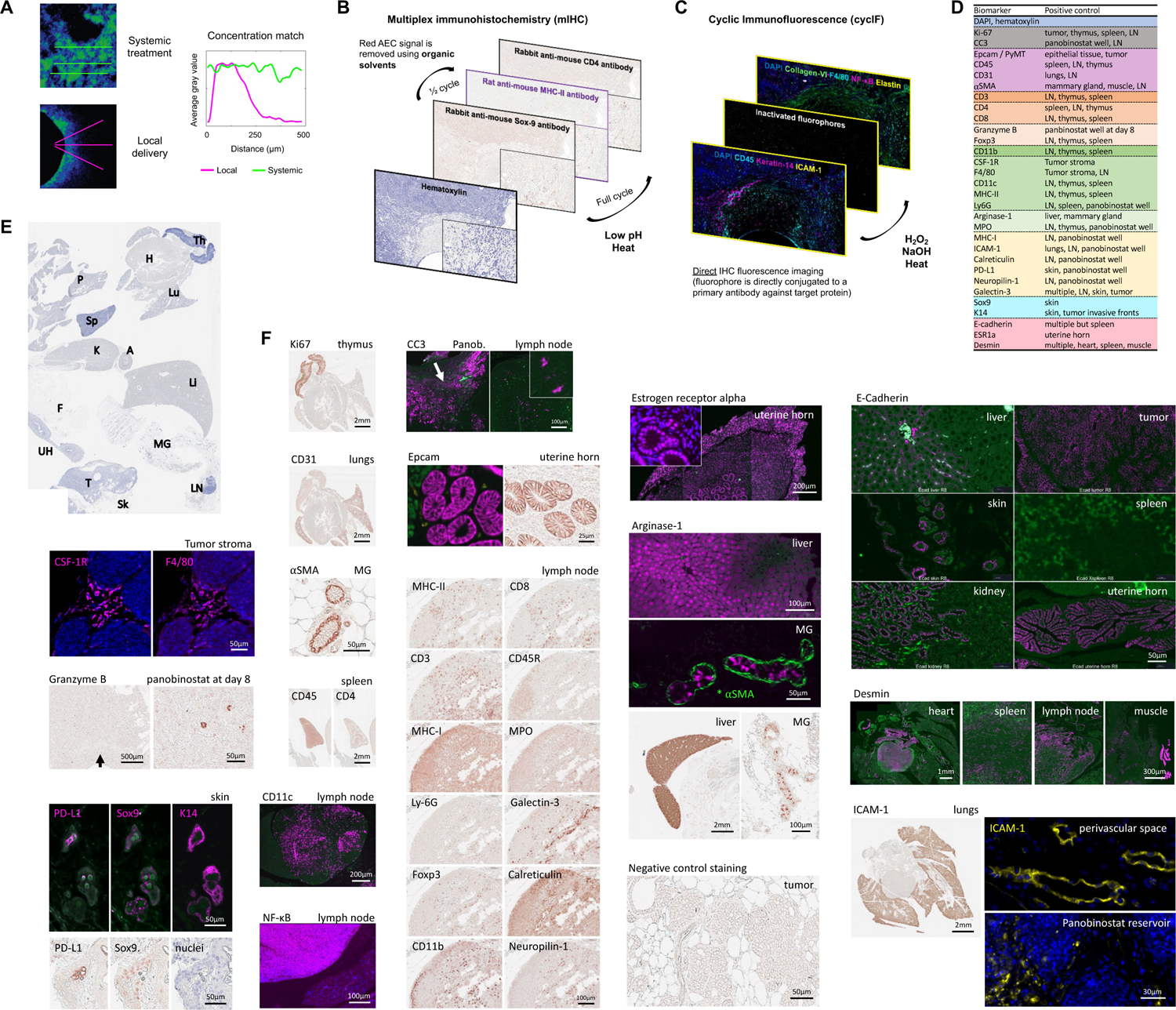
Components of the MIMA system and mIHC/cycIF anti-mouse antibody validation; Related to Figure 1. (A) Dosing for individual drugs was calibrated using mass spectrometry measurements comparing concentration of the same drug in situ after systemic treatment versus after local delivery (adapted and modified from (Jonas et al., 2015)). Sample images of intratumoral doxorubicin distribution at 6 hours after systemic treatment (top left image) and PEG-formulated doxorubicin transport from device at 20 hours after release (bottom left image). Signal mean intensity was extracted (n=3 each) and plotted using a moving average window filter to smoothen the signal (right). For detailed information on the pharmacokinetics of intratumoral drug release from the IMD see (Jonas et al., 2015). (B,C) Schematic overview of multiplex immunohistochemistry (mIHC; B) and cyclic immunofluorescence (cycIF, C). (C) mIHC utilizes indirect staining, iterative deposition of chromogen/enzyme pairs and brightfield microscopy to image the target signal. The chromogen used in this study is called 3-amino-9-ethylcarbazol (AEC) and it produces a red precipitate when visualized with polymer-based peroxidase conjugated to a secondary antibody (anti-rabbit or anti-rat; Histofine® Simple Stain). AEC is susceptible to organic solvents which is used to remove the red signal and detect two target proteins in one cycle. Primary antibody mixture is stripped in heated low pH citrate buffer is every cycle after scanning in order to further multiplex the staining on a single FFPE slide. Antibodies raised in rabbit and rat hosts alternate to prevent crosstalk between cycles. Hematoxylin counterstains nuclei to allow cell count and downstream image analysis (Figure S2). (C) cycIF utilizes fluorophores as reporters via direct labeling. Four to five non-overlapping fluorescent signals can be detected in a single cycle against dark background. DAPI signal is used to visualize nuclei for cell count. Progressive staining is enabled by inactivating the fluorophore using base hydrogen peroxide mixture and heat. Antibody specificity is cross-validated by performing chromogenic mIHC on the adjacent tissue section. (D) List of biomarkers (left column) and positive control tissues used for antibody validation and signal thresholding. (E) Hematoxylin staining of an FFPE section containing all positive control organs from an adult wild type FVB/N female mouse: thymus (Th), heart (H), lungs (Lu), liver (Li), mammary gland (MG), lymph node (LN), spleen (Sp), pancreas (P), adrenal gland (A), kidney (K), fat (F), uterine horn (UH). Tumor (T) with implanted device and attached skin (Sk) was embedded into the same FFPE block. (F) Representative images of individual biomarkers using mIHC (red signal in bright field images) or cycIF (magenta signal in fluorescent images unless otherwise stated). Biomarker name is located on the top left; while the name of the organ is located on the top right side of each image, respectively. Some positive signal can be detected in a macroscopic view (Ki67, CD31, CD4, CD45, NF-KB, desmin, arginase-1, ICAM-1). Section stained without primary antibody served as negative control in the mIHC procedure (last image). Green fluorescent channel served to detect autofluorescence and to separate background from specific staining in the cycIF procedure. Only antibodies with very strong specific staining such as aSMA (marked with a star *), were used in conjugation with Alexa fluor-488. Scale bars; shown.

**Figure S2.**
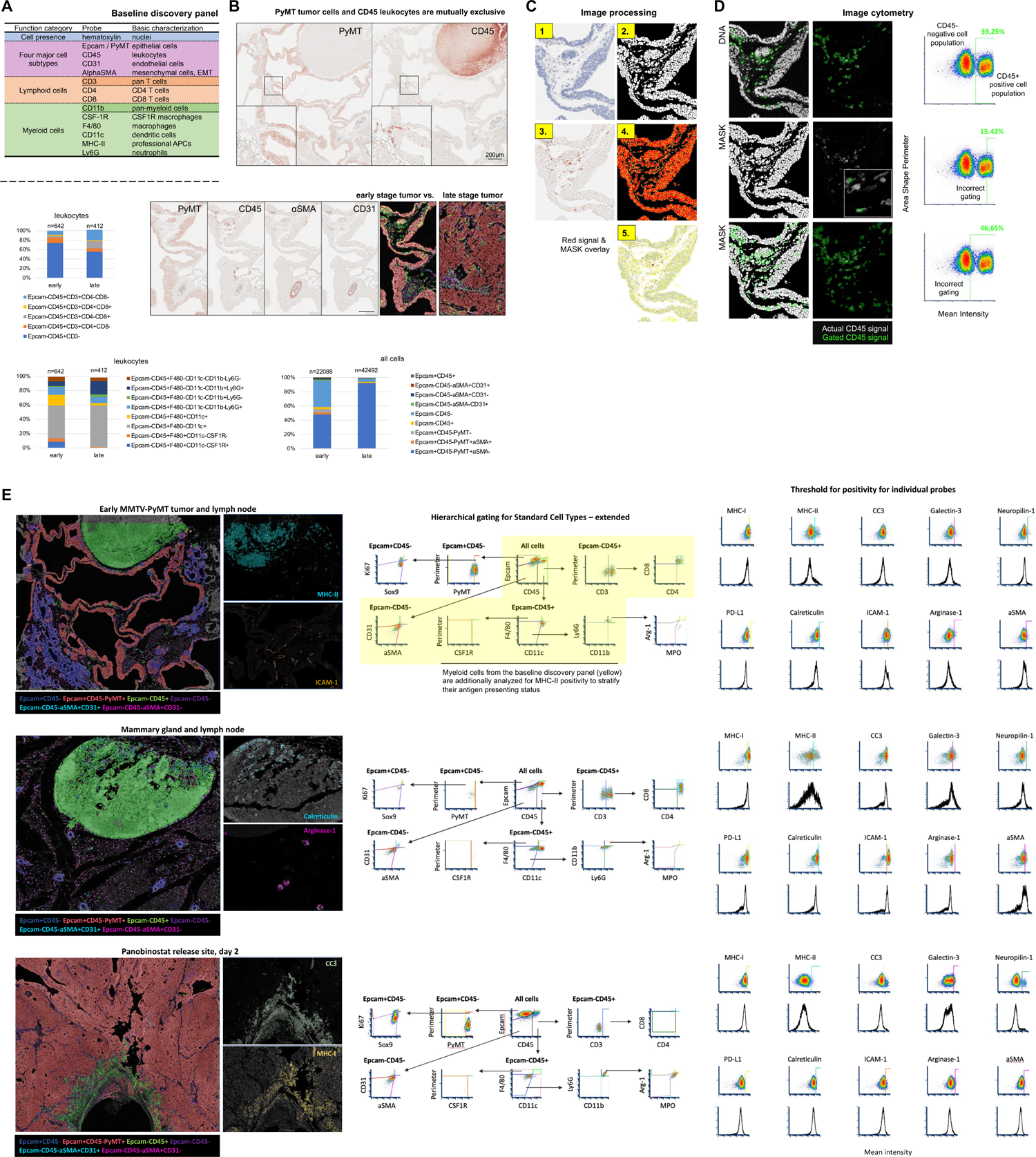
Analytical design to quantify single cell events in MIMA; Related to Figure 1. (A) The baseline discovery readout panel for MIMA is composed of a total of 13 probes we identified to be the minimal and satisfactory requirement to capture the major TME states to predict drug-induced changes which are actionable (see also Table S4). (B) Tissue section of an early MMTV-PyMT mammary carcinoma and adjacent lymph node (brightfield image of PyMT, CD45, *α*SMA, CD31 shown) was used to establish hierarchical gating strategies in image cytometry (in E) to define “standard cell types”. This for two reasons: presence of a lymph node in the same section offers the possibility to utilize mutual exclusivity (top) for reproducible signal thresholding. Second, early tumors provide with the opportunity to evaluate relatively broader range of phenotypically distinct cell types as compared to late-stage tumors (quantification, bottom). Number of cells analyzed is shown; data is derived from one and two tumors for early and late tumor sample, respectively. (C) Image processing for image cytometry analysis is composed of the following steps, briefly: hematoxylin staining (1) is color-deconvoluted and the signal is segmented using ImageJ watershed function (Schneider et al., 2012) to generate mask (2). Red AEC signal (3) mean intensity in a selection as defined by mask (4) is calculated for each cell (5). (D) Pixel intensity measurements and shape size measurements are used to gate cells for positive marker expression (CD45 in this case). FCS Express 6 and 7 Image Cytometry Software (De Novo Software), was used to obtain accurate thresholding using the cell population shape and dimensions. Correct gating is also monitored through visual inspection (second column). (E) Density plot of dimensionality reduction in hierarchical clustering to define “standard cell types” (middle column). The shape of the gates was designed to obtain quantitatively reproducible multiplex data, batch to batch, independent of the condition measured: early tumor and lymph node (top row), mammary gland and lymph node (middle row) and panobinostat implanted tumor sample two days post exposure (bottom row) are shown for comparison. For probes other than “standard cell types” (pleiotropic/undefined biology), threshold for positivity was determined manually using FCS Express 6 and 7 Image Cytometry software and positive control tissue (Figure S1D-F) (right). Sample pictures for marker positive cells; left.

**Figure S3.**
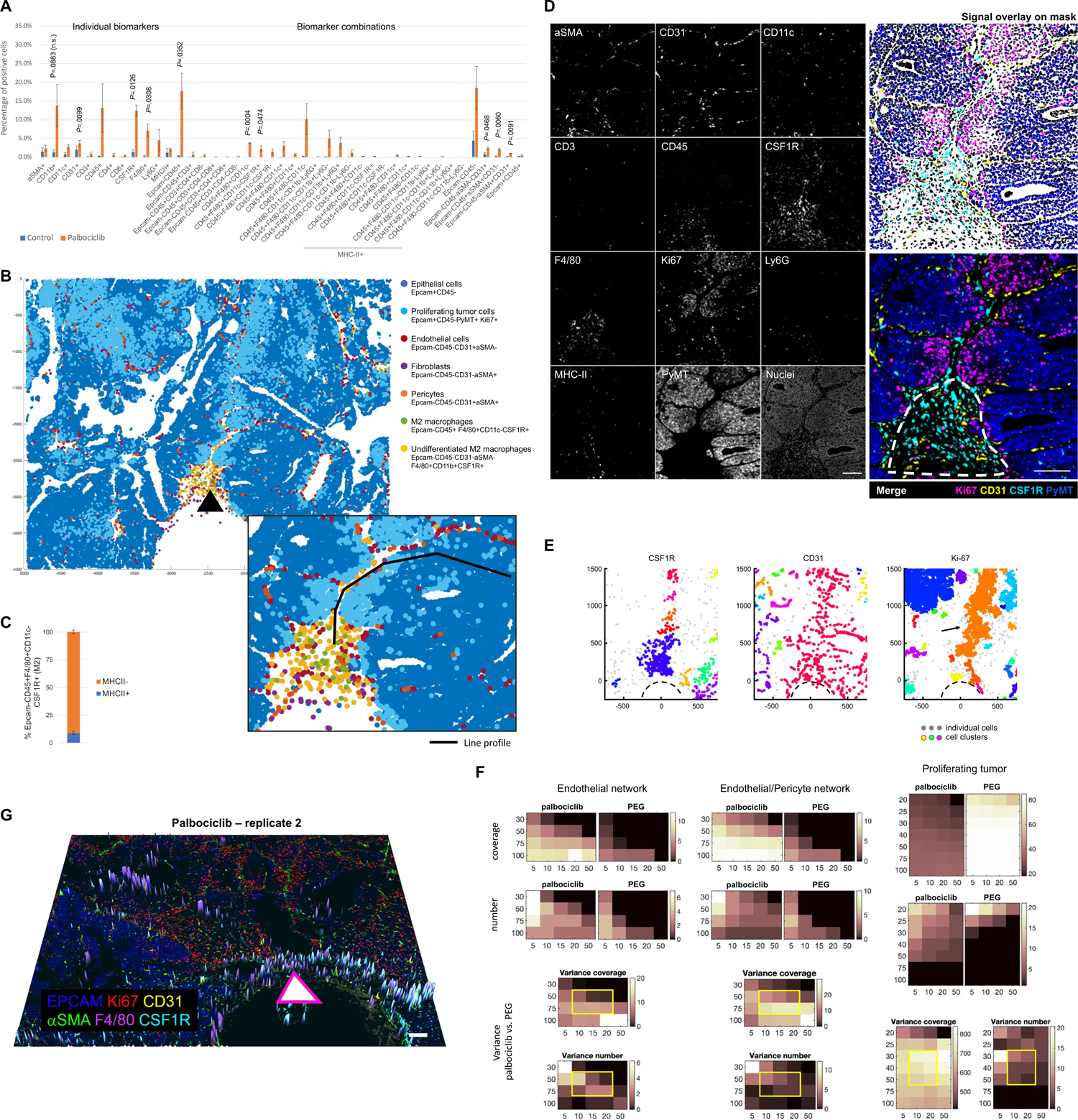
Locally induced tumor-TME changes at the palbociclib delivery sites; Related to Figure 2. (A) Quantification of single cell events using individual markers and marker combinations (including standard cell types). Bars are mean ± s.e.; n=3 palbociclib reservoirs in three tumors from three different MMTV-PyMT mice with late stage (d93-d100) spontaneously growing tumors implanted with IMD for three days. Significance was calculated by paired sample one tailed *t-*test. (B) Presentation of key response cell types (biomarker combination displayed) in XY space. Black arrow shows the drug releasing site; direction of the release is from the bottom to the top. The black line depicts region analyzed to quantify relative abundance of cell types with distance from the well in Figure 2D. (C) Percentage of MHC-II+ protumorigenic (M2) macrophages as defined by standard cell types. Stack bars are mean ± s.e.; n=3 palbociclib reservoirs in three tumors from three different mice. (D) Gray scale single channel images of depicted markers at the palbociclib reservoir (left) and merge composite images with or without overlay on the nuclei defined mask (top right and bottom right, respectively). Dashed line stratifies the “immediate pool” zone for Figure 2C. Scale bar, 100μm. (E) Distance based clustering of CSF1R, CD31 and Ki67 positive cells as a set of XY coordinates. Coordinate [0,0] identifies the drug source. Direction of the drug release is always from bottom to the top. Individual clusters were identified by minimum 10 cells within maximum distance 50μm, 75μm and 30μm for CSF1R+, CD31+ and Ki67+, respectively. Clusters were merged together if present within the maximum distance range. Each cluster is depicted with a different randomized color; individual (non-clustering) cells are shown as light gray points. Function was adapted and modified from Yann Marcon (Matlab, Oct 2019). Note larger cluster formation when analyzing individual markers as compared to standard cell types (Figure 2E) suggesting other than standard cell types express the CSF1R and CD31 marker (potential cell trans-differentiation). (F) Systematic testing of endothelial cell, endothelial and pericyte (union) cell and proliferating tumor cell cluster formation at palbociclib and control PEG reservoir based on cluster size presented in form of a heatmap. Cluster size was defined by minimal number of cells (x axis) within maximum distance range (y axis). Total coverage, number of clusters in the assay area variance between palbociclib and PEG in these two parameters was evaluated. Yellow rectangle defines cluster sizes that form specifically at the palbociclib stimulus site and have maximal variance (PEG vs palbociclib). Treatment specific cluster formation appears when minimum 10-20 cells are present with 50-75μm and 30-50μm for endothelial/pericyte cells and proliferating tumor cells, respectively. (G) Three-dimensional composite image of another palbociclib tumor tissue section. F4/80 macrophage marker is presented in high-view. Triangle arrow, which shows the localization and direction of the drug release, is shifted slightly to the right so that both normal tissue and Palbociclib affected region can be seen at once. Note slightly different extent of the TME response as compared to replicate number 1 (Figure 2B), while the shape and the order of the cell response with distance remains the same: CSF1R+, F4/80+ macrophages located in close proximity to the well; CD31 *α*SMA pericyte form network outside this region and Ki67 proliferating cells appear de novo (in the local microculture) around network.

**Figure S4.**
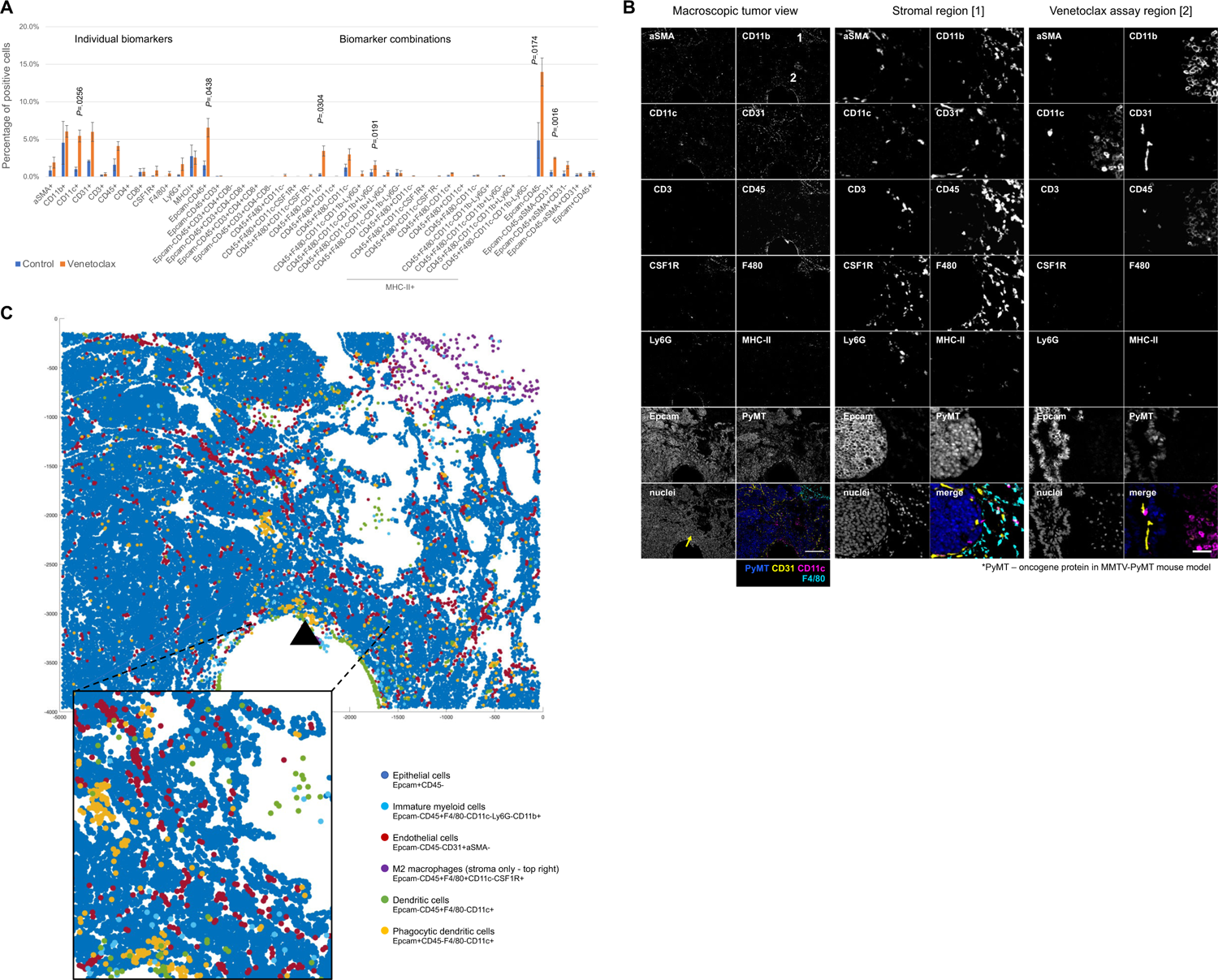
Locally induced tumor-TME changes at the venetoclax delivery sites; Related to Figure 3. (A) Quantification of single cell events at the venetoclax delivery site in spontaneous MMTV-PyMT tumors at three days of exposure by using individual markers and marker combinations (including standard cell types). Total cell counts to define percentage of positivity were between 3000 to 5000 cells per assay area and were matched ± 300 total cells for paired samples (experimental vs control region). Minimum population proportion within 5% margin of error and 95% confidence level was set to 0.75% (represents 12 cells) to discriminate noise from specific cell enrichment. Bars are mean ± s.e.; n=3 venetoclax reservoirs in two tumors from two different mice. Significance was calculated by paired sample one tailed t-test. (B) Gray scale single channel images of depicted markers at the Venetoclax reservoir. Macroscopic view is on the left; magnified view of stromal regions and venetoclax proximal region are in the middle and right, respectively. Composite of depicted markers is in the colored image. Scale bar, 500μm and 50μm for macroscopic and zoomed view, respectively. (C) Presentation of key response cell types/states (biomarker combination displayed) in XY space. [0,0] coordinate is the drug releasing site; direction of the release is from the bottom to the top.

**Figure S5.**
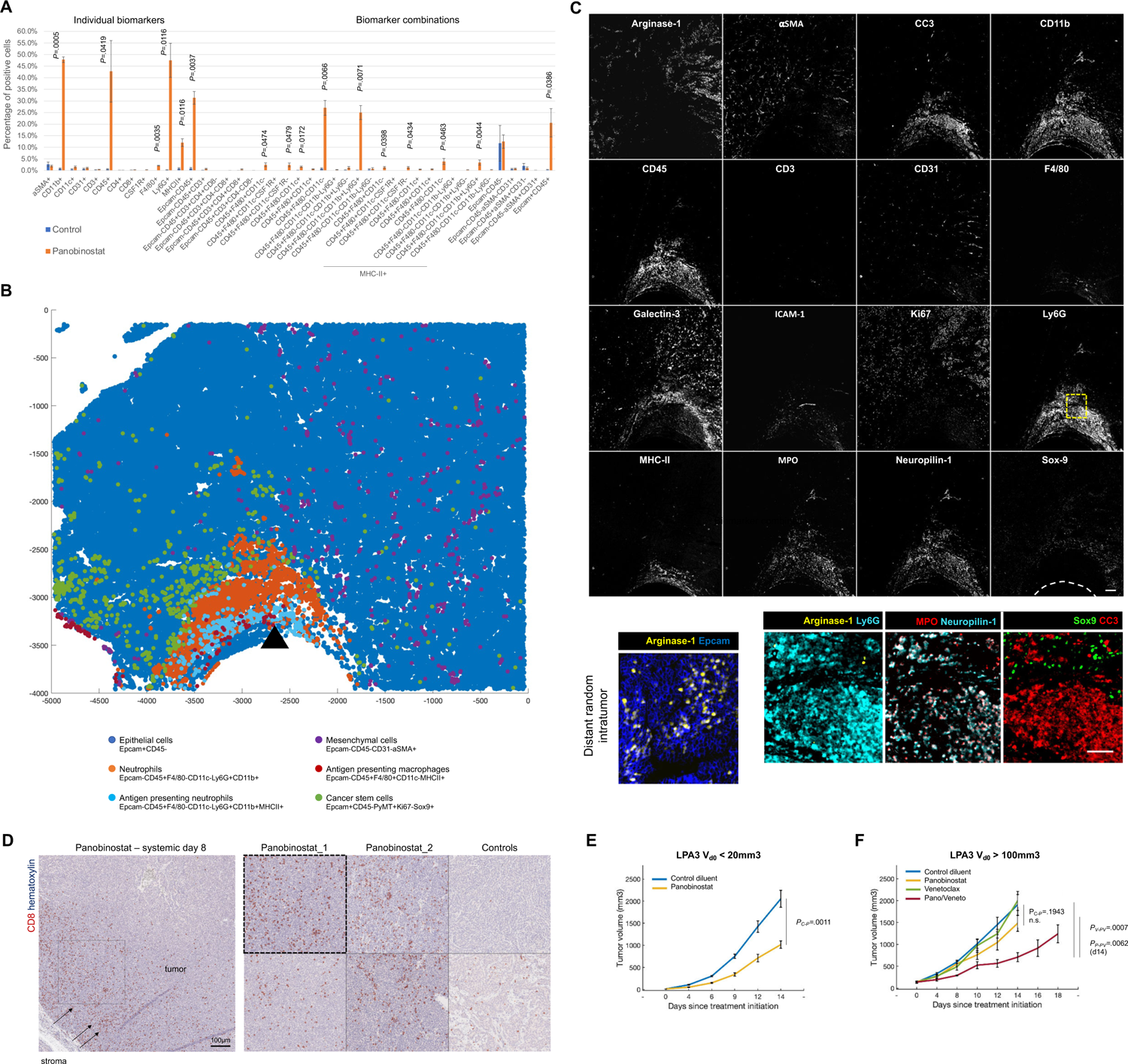
Local and systemic effects of the pan-HDAC inhibitor, panobinostat, in different mouse models of breast cancer. Related to Figure 4. (A) Quantification of single cell events at the panobinostat delivery site in spontaneous MMTV-PyMT tumors at three days of exposure by using individual markers and marker combinations including standard cell types. Total cell counts to define percentage of positivity were between 3000 to 5000 cells per assay area and were matched ± 300 total cells for paired samples (experimental vs control region). Minimum population proportion within 5% margin of error and 95% confidence level was set to 0.75% (represents 12 cells) to discriminate noise from specific cell enrichment. Bars are mean ± s.e.; n=3 panobinostat reservoirs in two tumors from two different mice. Significance was calculated by paired sample one tailed t-test. (B) Presentation of the most prominent response cell types (biomarker combination displayed) in XY space. Black arrow marks source and direction of drug release. (C) Gray-scale single channel images of depicted markers at the panobinostat reservoir (replicate 1). Dashed line in the Sox9 image marks the border of the device. Scale bar, 100μm. Dashed yellow box in the Ly6G image marks magnified view of a region at the intersection of dying cells (by CC3) and surrounding TME. Scale bar, 50μm. (D) ErbB2*Δ*Ex16 mice with spontaneously growing tumors were treated with diluent (control) and panobinostat systemically for 4 days, after which the tumors were extracted at day 7 and formalin fixed paraffin processed (FFPE). Images show tumor tissue sections stained with anti-mouse CD8 antibody (red AEC signal) and hematoxylin (blue). Note the gradient of high CD8 infiltration closer to stroma with decreasing tendency towards the tumor center (arrows) suggesting CD8 T cells are recruited from stroma regions. Scale bar, 100μm (E, F) Tumor growth rate in syngeneic LPA3 mice in which tumors were induced by orthotopic injection of primary tumor cells into mammary fat pad. The mice were treated systemically by depicted treatments as shown in the graph. Early (<20mm3; D) and more advanced (>100mm3, E) tumors were tested as for the treatment start (day 0). Mice were treated intraperitoneally with dose and schedule as defined in Figure 3G and 4H. Line graphs show mean ± s.e., n=5 tumors in five mice per group. Significance was calculated by two sample two-tailed t-test with equal variance.

**Figure S6.**
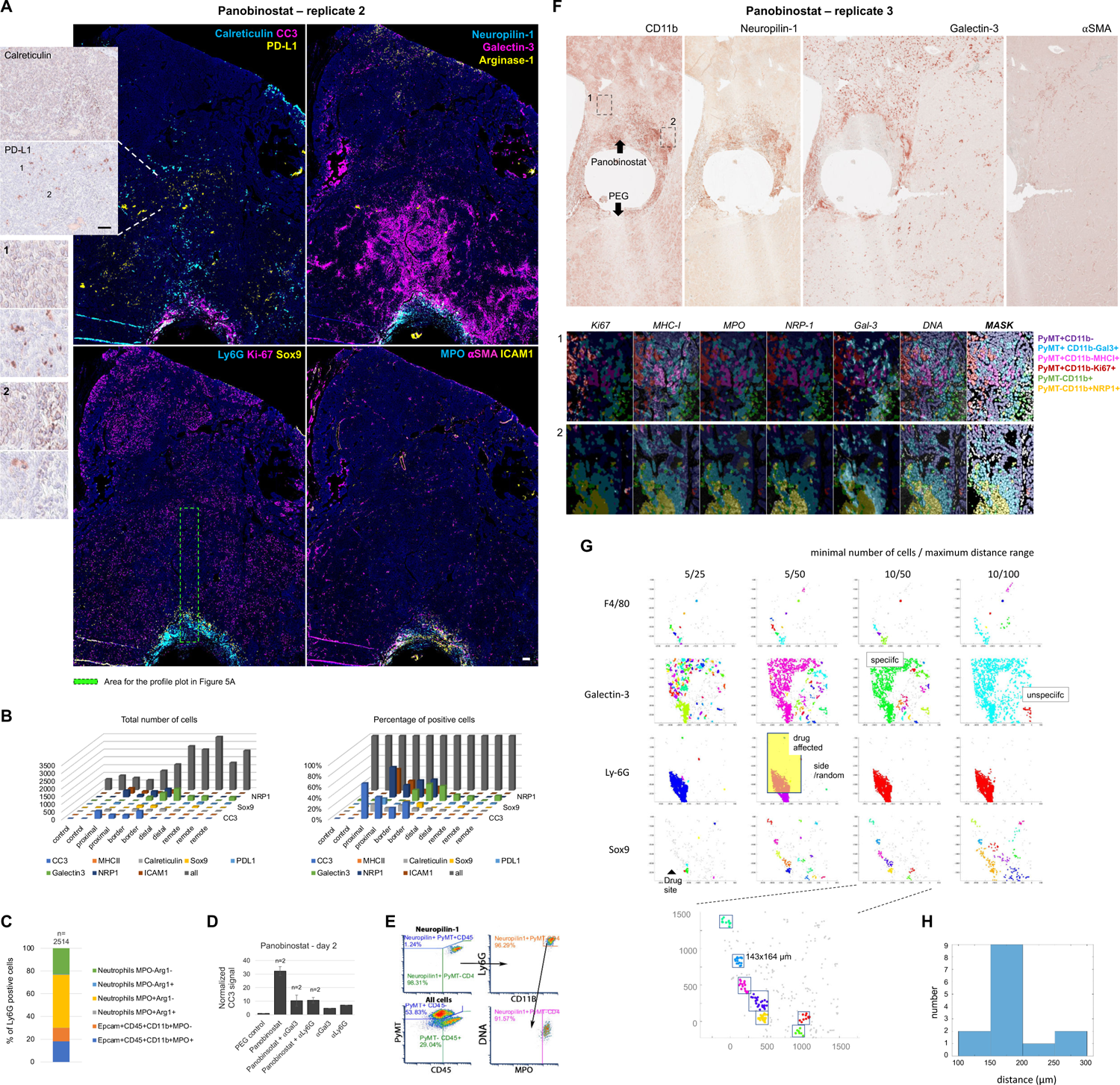
Biomarkers of immunogenic cell death and associated mechanisms of resistance induced by local panobinostat drug stimulus; Related to Figure 5. (A) Large field of view three-color composite images showing biomarkers of immunogenic cell death induced by panobinostat (replicate 2) reservoir at three days of exposure. Calreticulin and PD-L1 IHC (red AEC signal) overlayed on hematoxylin nuclei (in blue) are presented in bright field zoomed image on the left. (B) Quantification of single cells positive cells for depicted biomarker with distance from the reservoir; total cell counts (left) and rate of positive cells (right) are presented in form of a 3D bar graph. 2-3 ROIs are presented per assay zone (proximal, distal, distal, remote, control). (C) Expression rate of CD45, MPO and arginase-1 on Ly6G+ cells in the panobinostat assay area to stratify phagocytic, cytotoxic and immune suppressive neutrophils, respectively. Number (n) of analyzed cells is presented. (D) Panobinostat reservoirs were co-loaded with anti-Ly6G (clone 1A8) and galectin-3 (clone M3/38) antibodies at 5:1 to 10:1 ratio and CC3 IHC signal was quantified at the drug releasing site. n=2 for experimental and 1 for control conditions, respectively. All results were obtained from a single IMD in one tumor which was implanted for two instead of typical three days to account for antibody half-live. (E) Image cytometry measuring neuropilin-1 expression on cytotoxic neutrophils. For comparison, population distribution of all cells is presented on the bottom left. (F) Bright field large field of view images of CD11b, neuropilin-1, galectin-3 and *α*SMA IHC at the panobinostat reservoir (replicate 3) at 3 days of exposure. Zoomed images show color-coded extracted signal overlayed on the true signal of the depicted biomarkers (white). (G) Distance based cluster analysis testing different cluster size strategies to identify treatment specific cluster formation. The function implements a user defined cluster sizes set by minimal number of cells (first number in the top right legend) within maximum distance range (second number in the top right legend) to display cluster formation in randomized color while individual cells not meeting the clustering criteria are presented in gray. Drug source is shifted to the left to stratify cluster formation in assay area (proximity to the well) vs side/random regions. Clustering strategies using low cell number (e.g. 5 cells, first two columns) and high distances (e.g. 100μm, right column) show clusters forming unspecifically outside the assay area; Clustering strategy using minimal 10 cells within maximum distance range 50μm (10/50 column) shows cluster formation specifically above the drug site for all presented markers (F4/80, Galectin-3, Ly6G, Sox9). Magnified Sox9 cluster formation; bottom. (H) Frequency of Sox9 cluster sizes. Cluster size around 175µm in diameter, which were the most prominent, were used for downstream analysis of pairwise proximity measurements of Sox9 with other markers (Figure 5G, H).

**Figure S7.**
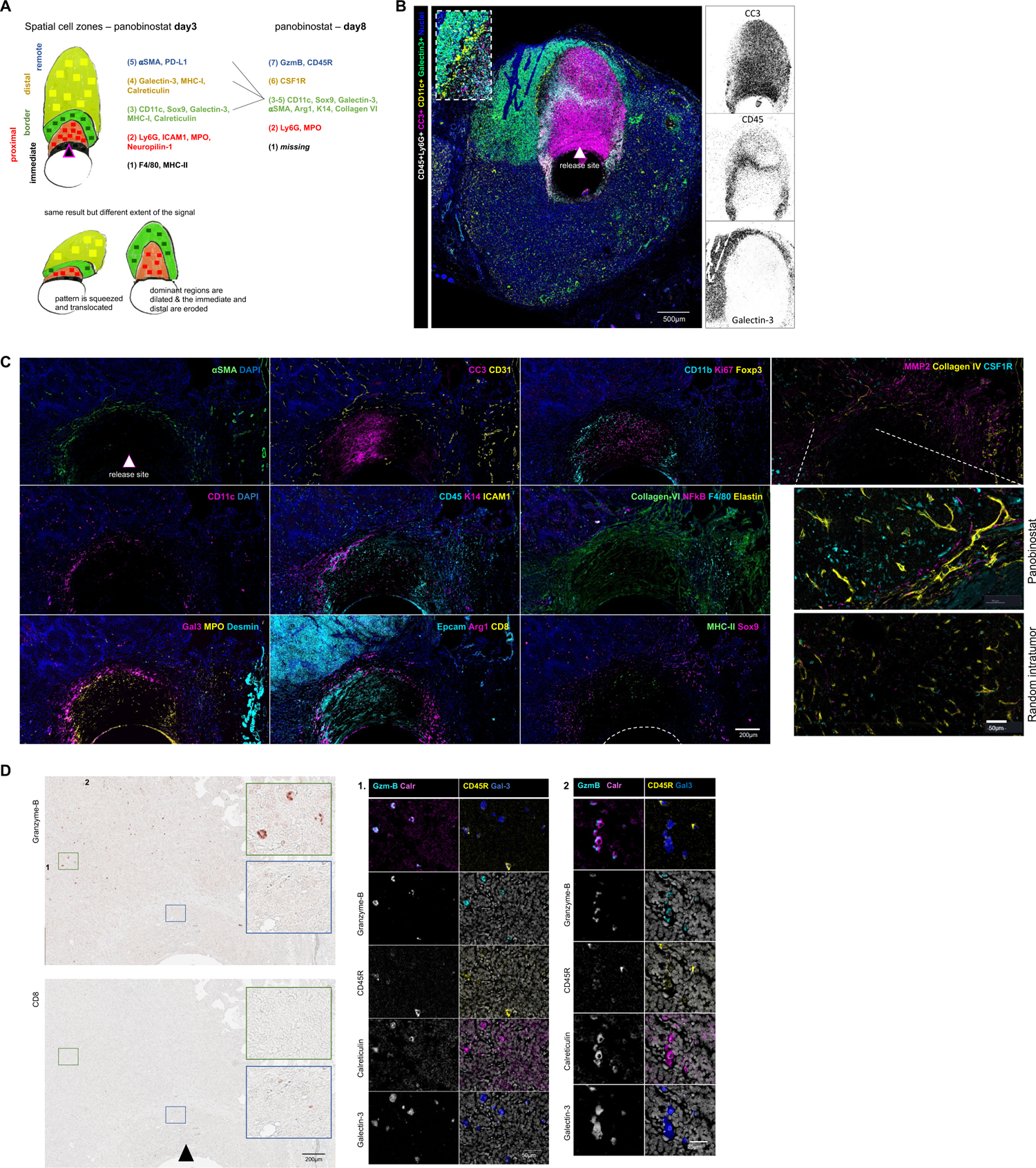
Local panobinostat efficacy in other mouse model and at a later (day 8) timepoint; Related to Figure 5. (A) A schematic presentation of cell phenotype separation into zones with distance from the panobinostat reservoir at day three and day eight timepoint. Shared phenotypes between the two timepoints suggest order of the cell transition with distance from the well defines the sequence of cellular events (earliest to latest) and is as follows: 1) MHC-II antigen presenting cells and F4/80 macrophages are recruted first to the drug delivery well as they are located *immediately* at the drug well at early timepoint; 2) MPO cytotoxic, ICAM1 adhesive/activated, neuropilin-1 positive N1 neutrophils are recruited second (*proximal* zone) and this phenotype is halt by the *border* barrier composed of 3) CD11c dendritic cells, Sox9 cancer stem cells and galectin-3. Relative increase of MHC-I and calreticulin on the cell surface starts to form in this border zone and propagates to (4) *distal* region with decreasing gradient profile. 5) galectin-3 expression is associated with PD-L1 and is halt by aSMA fibroblast *remotely* from the well. Over time, the last three zones (3-5) merge into a single border zone composed of CD11c dendritic cells, arg-1 immune suppressive cells, Sox9 cancer stem cells, K14 cells of invasive front and ECM deposition/processing componenets (collagen VI and MMP2). The immediate macrophage zone is missing at the day eight timepoint; instead, two new cell types appear at the new border zone: (6) CSF1R+ cells (C) and (7) granzyme B cytotoxic CD45R B cells (D). (B) ErbB2*Δ*Ex16 mice with spontaneously growing tumors were implanted with IMD loaded with panobinostat and the tumor with the device in place was extracted at three days and was formalin fixed and paraffin processed. Picture shows a five-color composite image of biomarkers associated with immunogenic cell death. While the absolute extent of the phenotype is larger as compared to those observation in MMTV-PyMT model; the order of the phenotypes (spatial cell pattern with distance from the drug source) remains identical with Ly6G+ leukocytes and CC3 apoptosis present at the proximal, CD11c dendritic cells present at border and galectin-3 present at the distal regions. We note that the ErbB2*Δ*Ex16 model of breast cancer express both basal and luminal cytokeratin markers (Turpin et al., 2016). The different extent of the signal might be associated with the tumor subtype difference or the compactness/”fluidity” of the tumor tissue. (C) Sectioned tissue surrounding the implantable microdevice containing panobinostat for eight days was stained with cycIF using panel of mouse specific antibodies (Table S3) to display cellular phenotypes supporting the panobinostat model of response describe in (A). (D) Adjacent section as described in (C) was stained by multiplex immunohistochemistry using mouse specific antibodies (Table S2) and displays presence of cytotoxic granzyme B+ CD45R+ B cells expressing calreticulin and galectin-3. No other marker was expressed on these cells (or under the IHC sensitivity limit; not shown). Also, no other cells expressed the granzyme B cytotoxicity marker intratumorally; inside the tumor bed. Panobinostat drug release site is marked by triangle in all images. Direction of the drug release is always from the bottom to the top. Scale bars; shown.

**Figure S8.**
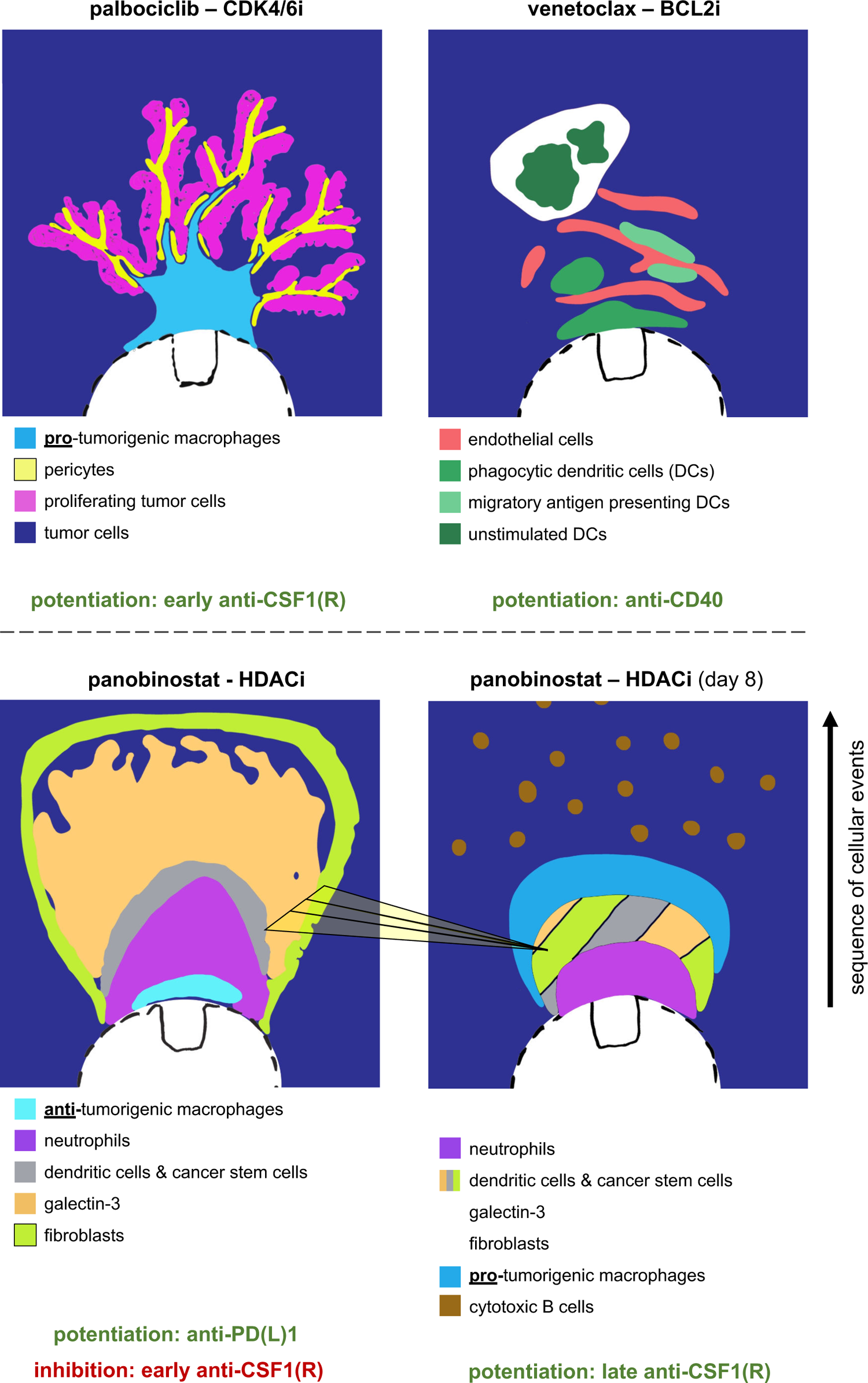
Illustration of complex tumor tissue response to targeted therapies and rational for immune modulatory combinations. Schematic presentation of the spatial cell associations induced by palbociclib, venetoclax and panobinostat at three days of local drug exposure; and panobinostat at eight days of drug exposure (bottom right). Palbociclib efficacy is associated with a “tree” or “delta-like” cell organization with targetable protumorigenic macrophages being localized in immediate proximity to the reservoir leading to endothelial/pericyte network formation and proliferation of tumor cells in proximity to this network. The optimal schedule of immunotherapy application can be estimated based on the location of the targetable phenotype from the drug well. Early (pre-treatment) modulation of protumorigenic macrophages using CSF1/CSF1R targeted immunotherapy was used to potentiate the efficacy of palbociclib. Venetoclax induces small vessel formation and recruitment of dendritic cells which appeared as “split clusters” at the drug well and it is yet unknown whether and how the subsets are functionally related. Anti-CD40 mediated licensing of these DCs can shift the balance from immune tolerance to T cell priming and this immunotherapy was applied at three days of venetoclax treatment since DCs were present in all regions of the assay area. Panobinostat induces immunogenic cell death associated with infiltration of antigen presenting neutrophils and macrophages but the propagation of this positive response is limited by enrichment of cancer stem cells and recruitment of dendritic cells and fibroblasts which merge into a resistant barrier over time and subsequently induce recruitment of protumorigenic macrophages and cytotoxic B cells. The cellular pattern of response has a “layering” or bay-like” structure implying more intense cell interaction might be involved at the cell layer interfaces. Anti-PD-1 immunotherapy was identified as the rational combination partner for panobinostat due to induction of immunogenic cell death and increased antigenicity. While early (pretreatment) administration of CSF1/CSF1R targeted immunotherapy can negatively affect antitumor function of macrophages (tested, Figure 4C); late administration of this immunotherapy might be beneficial to deplete/polarize the protumor macrophage which are recruited at the later timepoint (hypothetical).

**Figure S9.**
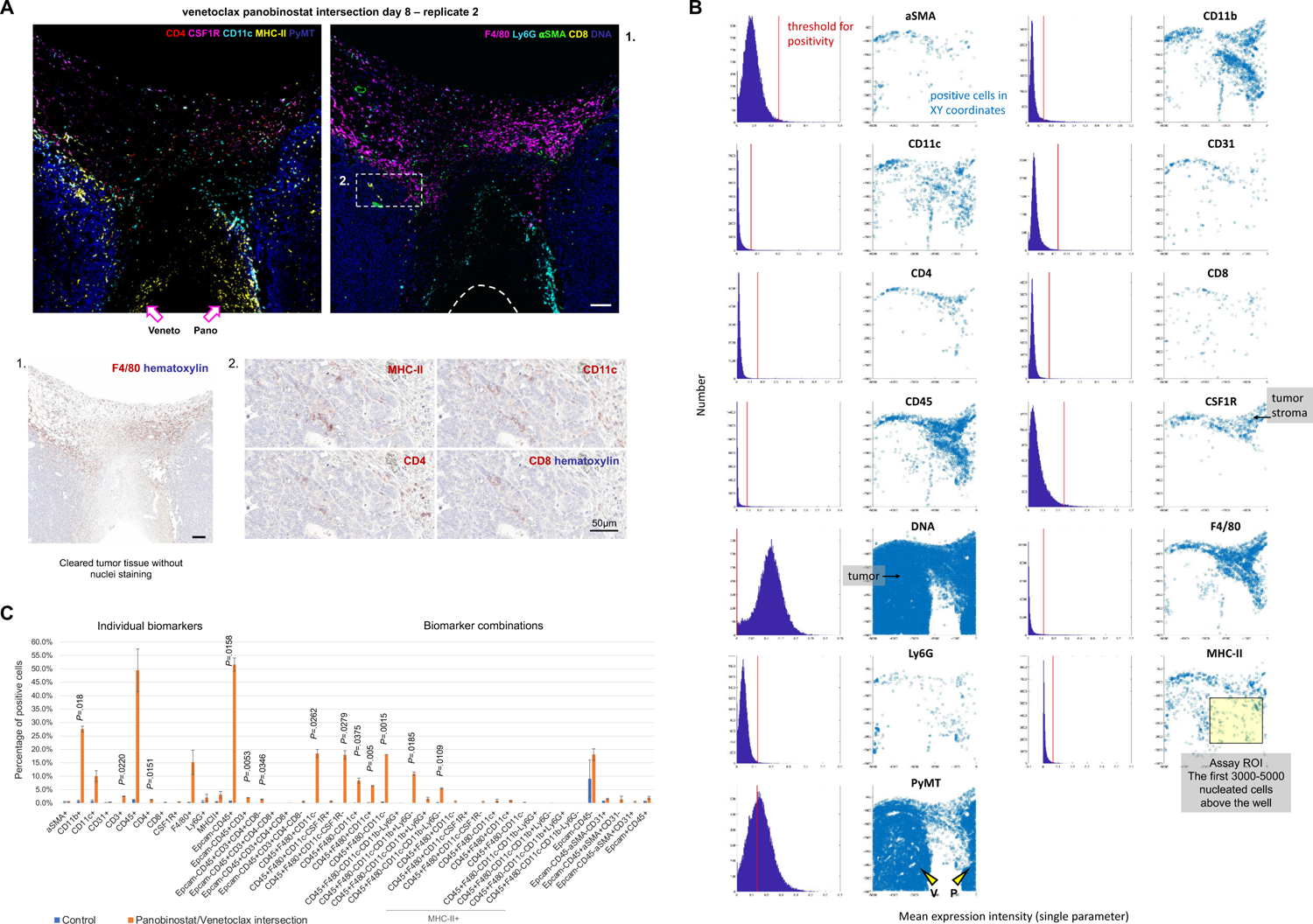
Local and systemic effects of the pan-HDAC inhibitor, panobinostat, in different mouse models of breast cancer. (A) Quantification of single cell events at the intersection of venetocalx and panobinostat delivery sites in orthotopic MMTV-PyMT tumors at eight days of exposure by using individual markers and marker combinations. Total cell counts to define percentage of positivity were between 3000 to 5000 cells per assay area and were matched ± 300 total cells for paired samples (experimental vs control region). Minimum population proportion within 5% margin of error and 95% confidence level was set to 0.75% (represents 12 cells) to discriminate noise from specific cell enrichment. Bars are mean ± s.e.; n=2 wells from one tumor in one mouse. Significance was calculated by paired sample one tailed t-test. (B) Macroscopic view of marker positive cells at the panobinostat/venetoclax drug intersection. Histograms show mean expression intensity of individual biomarkers. Red line defines threshold for positivity. Marker positive cells as defined by threshold are presented as blue dots in XY space. Black arrow is pointing on the tumor stroma region remote from the reservoir. Yellow triangle arrows indicate the source and direction of the drug release. Yellow box shows the approximation of the assay area. (C) Five-color composite images showing most prominent biomarkers induced at the intersection of panobinostat and venetoclax. White arrows indicate the source and direction of the drug release. Scale bar; 200μm. Macroscopic view of tumor tissue sections stained with anti-mouse F4/80 (red AEC signal) and hematoxylin (blue). Tumor cleared tissue is shown by lack of hematoxylin staining in the center suggesting lack of nucleated cells (bottom left image). Magnified view of intratumoral T cell infiltration to professional antigen presenting DC region near panobinostat/venetoclax assay area. The device was implanted for eight days in MMTV-PyMT tumor induced by orthotopic implant into mammary fat pad of syngeneic mice.

